# Correlative light and electron microscopy reveals that mutant huntingtin dysregulates the endolysosomal pathway in presymptomatic Huntington’s disease

**DOI:** 10.1101/2021.03.05.433899

**Authors:** Ya Zhou, Thomas R. Peskett, Christian Landles, John B. Warner, Kirupa Sathasivam, Edward J. Smith, Shu Chen, Ronald Wetzel, Hilal A. Lashuel, Gillian P. Bates, Helen R. Saibil

## Abstract

Huntington’s disease (HD) is a late onset, inherited neurodegenerative disorder for which early pathogenic events remain poorly understood. Here we show that mutant exon 1 HTT proteins are recruited to a subset of cytoplasmic aggregates in the cell bodies of neurons in brain sections from presymptomatic HD, but not wild-type, mice. This occurred in a disease stage and polyglutamine-length dependent manner. We successfully adapted a high-resolution correlative light and electron microscopy methodology, originally developed for mammalian and yeast cells, to allow us to correlate light microscopy and electron microscopy images on the same brain section within an accuracy of 100 nm. Using this approach, we identified these recruitment sites as single membrane bound, vesicle-rich endolysosomal organelles, specifically as (i) multivesicular bodies (MVBs), or amphisomes and (ii) autolysosomes or residual bodies. The organelles were often found in close proximity to phagophore-like structures. Immunogold labeling localized mutant HTT to non-fibrillar, electron lucent structures within the lumen of these organelles. In presymptomatic HD, the recruitment organelles were predominantly MVBs/amphisomes, whereas in late-stage HD, there were more autolysosomes or residual bodies. Electron tomograms indicated the fusion of small vesicles with the vacuole within the lumen, suggesting that MVBs develop into residual bodies. We found that markers of MVB-related exocytosis were depleted in presymptomatic mice and throughout the disease course. This suggests that endolysosomal homeostasis has moved away from exocytosis toward lysosome fusion and degradation, in response to the need to clear the chronically aggregating mutant HTT protein, and that this occurs at an early stage in HD pathogenesis.

## Introduction

Huntington’s disease (HD) is a late onset, inherited neurodegenerative disorder that manifests with psychiatric disturbances, movement disorders and cognitive decline [4]. It is caused by a CAG repeat expansion in exon 1 of the huntingtin gene (*HTT*) that encodes an abnormally long polyglutamine (polyQ) tract in the huntingtin protein (HTT) [25]. Individuals with CAG expansions of 35 or less remain unaffected, those with 40 or more will develop HD within a normal lifespan, and 36-39 CAGs confer an increasing risk of developing the disease [62]. CAG expansions of approximately 65 or more can result in onset during childhood or adolescence [70]. Huntingtin aggregates are deposited in the brains of HD patients in the form of inclusions that are predominantly cytoplasmic and scattered throughout the neuropil in cell bodies, dendrites and axons [11,18]. They also occur as nuclear inclusions, the frequency of which increases with increasing CAG repeat length [11]. Neuronal cell death occurs in the striatum, cortex and other brain regions [50,74].

The propensity for mutant HTT to aggregate is governed by the extent of the polyQ expansion, and the length of the N-terminal HTT fragment in which it resides [66]. The exon 1 HTT protein is particularly aggregation-prone [66]. An incomplete splicing event that occurs in knockin mouse models of HD, as well as patient brains and cell lines, results in a small polyadenylated transcript that encodes this highly pathogenic protein [65,51]. In addition to this, N-terminal fragments of mutant HTT are generated by the proteolysis of full-length HTT [16,29,36,77]. Exon 1 HTT self-assembles *in vitro* to form insoluble end-stage amyloid-based fibrillar aggregates with a cross-β-sheet architecture [63,67]. The initiating exon 1 HTT associations are thought to occur through α-helix-rich interactions of the first 17 amino acids (N17), leading initially to spherical, α-helix rich oligomeric structures [71,26,57]. This may enhance the elongation of a stochastically-formed polyQ β-hairpin, facilitating nucleation of a β-sheet-based amyloid structural motif [21,78] to which additional exon 1 HTT monomers are recruited [75]. Our recent description of a liquid to solid phase transition for exon 1 HTT proteins indicates that the mutant HTT aggregation process in cells may involve liquid-like assemblies prior to the formation of fibrillar structures or other stable aggregates [56]. Regardless of the specific steps preceding nucleation, post-nucleation amyloid growth appears to proceed via oligomeric nanofibrils that possess key functional attributes of amyloid substructure: the ability to grow into mature fibrils by recruitment of monomeric exon 1 HTT [12,78].

HTT inclusions have been shown to have a fibrillar sub-structure *in vivo* and cryo-electron tomography of exon 1 HTT inclusions in cell models also revealed a fibrillar structure that distorted the organization of the endoplasmic reticulum (ER) and perturbed ER membrane dynamics [5,61,66]. Inclusions *in vivo* are complex structures which sequester many other proteins [24] and may perturb multiple processes critical to cellular function. Cell culture experiments previously suggested that the formation of inclusions might be part of a homeostatic protective mechanism [2]. However, subsequent experiments suggest that inclusions can be directly cytotoxic [44]. A recent study showed that an aggregated form of exon 1 HTT, possessing a fundamental amyloid substructure, was the toxic species of mutant HTT in HD flies and transfected neurons, although the exact nature of this amyloid species, i.e. an inclusion, dispersed fibril cluster, or nanofibril, was not determined [12,78]. The use of a seeding assay has demonstrated that minute quantities of small self-propagating exon 1 HTT fibrils were sufficient to dramatically shorten the lifespan of HD transgenic flies [3].

It was previously shown that a subset of cytoplasmic inclusion bodies in tissue sections from HD mouse models and HD patient *post-mortem* brains, termed “aggregation foci”, were competent to recruit polyQ peptides [52]. Here, we set out to determine the ultrastructure of these recruitment foci, which appeared indistinguishable from nuclear and other cytoplasmic inclusions by anti-HTT antibody staining. We found that fluorescently labelled exon 1 HTT proteins were recruited to sites within neuronal cell bodies, in brain sections from two distinct HD mouse models, in a polyQ-length and disease-stage dependent manner. In order to identify the structures of these recruitment foci, we established a correlative light and electron microscopy (CLEM) procedure to co-locate fluorescent signals with EM images on the same brain section. We found that the exon 1 HTT proteins were recruited to single membrane-bound, vesicle-rich organelles that resembled multivesicular bodies (MVBs) or amphisomes, whereas others, with single or multiple vacuoles, resembled autolysosomes or residual bodies. Immunogold labelling localized the mutant HTT to non-fibrillar, electron-lucent regions within the lumen of these organelles. Our finding that markers of MVB-related exocytosis were depleted in presymptomatic HD mice demonstrates that endolysosomal homeostasis is dysregulated at an early stage in HD pathogenesis.

## Materials and methods

### Animals

All procedures were performed in accordance with the Animals (Scientific Procedures) Act 1986 and were approved by the University College London Ethical Review Process Committee. zQ175 knock-in mice were generated by replacing exon 1 of mouse *Htt* with exon 1 from human *HTT,* carrying a highly expanded CAG repeat [20,49] from which the neo selectable marker has been removed (delta neo) [15] and were bred by backcrossing zQ175 males to C57BL/6J females (Charles River). R6/2 mice are transgenic for the 5’ region of the mutant human huntingtin *(HTT)* gene [47] and were bred by crossing R6/2 males to C57BL/6JOlaHsd x CBA/CaOlaHsd F1 females (B6CBAF1/OlaHsd, Envigo, Netherlands).

Within each colony, genetically modified and wild-type mice were group housed with up to five mice per cage, dependent on gender, but genotypes were mixed. Mice were housed in individually ventilated cages with Aspen Chips 4 Premium bedding (Datesand) with environmental enrichment which included chew sticks and play tunnel (Datesand). They had unrestricted access to food (Teklad global 18% protein diet, Envigo) and water. The temperature was regulated at 21°C ± 1°C and animals were kept on a 12 h light / dark cycle. The animal facility was barrier-maintained and quarterly non-sacrificial FELASA screens found no evidence of pathogens. For molecular analyses, mice were sacrificed by a schedule 1 procedure, brains were rapidly dissected, tissues were snap frozen in liquid nitrogen and stored at −80°C.

### Genotyping and CAG repeat sizing

Genotyping and CAG repeat sizing were performed by PCR of ear-biopsy DNA and all primers were from Invitrogen. For zQ175 mice, a 20 μL genotyping reaction contained 10 - 15 ng DNA, GoTaq Flexi buffer, 2.5 mM MgCl_2_, 0.2 mM dNTPs, 1.0 μM 19Fhum [5’-AGGAGCCGCTGCACCGA-3’], 1.0 μM 431R2 [5’-CTCTTCACAACAGTCATGTGCG-3’] and 1.0 U GoTaq2 polymerase (Promega). Cycling conditions were 30 s at 98°C, 34 x (15 s at 98°C, 15 s at 64°C, 30 s at 72°C), 5 min at 72°C. The amplified products were 324 bp for wild-type and 240 bp for zQ175. For R6/2 mice, a 10 μL reaction contained 50 - 100 ng DNA, 1.0 μM forward primer [5’-CGCAGGCTAGGGCTGTCAATCATGCT-3’], 1.0 μM reverse primer [5’-TCATCAGCTTTTCCAGGGTCGCCAT-3’], 1.0 μM forward internal control primer [5’-AGCCCTACACTAGTGTGTGTTACACA-3’], 1.0 μM reverse internal control primer [5’-CTTGTTGAGAACAAACTCCTGCAGCT-3’] and Dream Taq Hot Start Green PCR Master Mix (Thermo Fisher Scientific). Cycling conditions were 3 min at 95°C, 34 x (30 s at 95°C, 30 s at 60°C, 1 min at 72°C), 15 min at 72°C.

For CAG repeat sizing for both zQ175 and R6/2, a 25 μL reaction contained 25 - 50 ng DNA, 0.2 μM FAM-labelled forward primer [5’-CCTTCGAGTCCCTCAAGTCCTT-3’], 0.2 μM reverse primer [5’-CGGCTGAGGCAGCAGCGGCTGT-3’], AmpliTaq Gold 360 Master Mix (Thermo Fisher Scientific) and GC enhancer (Thermo Fisher Scientific). Cycling conditions were 10 min at 95°C, 34 x (30 s at 95°C, 30 s at 63°C and 90 s at 72°C), 7 min at 72°C. Samples were run on an ABI3730XL Genetic Analyser with MapMarker ROX 1000 (Bioventures) internal size standards and analyzed using GeneMapper v5 software (Thermo Fisher Scientific). The mean CAG repeat size for all zQ175 mice was 191.7 ± 4.6 (S.D.) and for R6/2 was 182.1 ± 1.8 (SD).

### Recruitment assay with biotinylated peptides

The recruitment of the mbPEGQ30 peptide and subsequent immunohistochemistry was performed as described in [52] with modifications. In brief, mice were perfused and 30 μm free-floating brain sections prepared as described previously [35]. Sections were washed in phosphate buffered saline (PBS) and treated with 1% sodium borohydride in PBS for 30 min. Borohydride was removed by washing three times in PBS. Sections were permeabilized and blocked by washing three times in PBS with 0.3% triton (PBST) and incubated overnight with 10 nM mbPEGQ30 in PBST at 4°C. Rigorous washing was performed to remove unbound peptides with PBST over several hours at room temperature. Sections were then washed three times in PBS before signal amplification using Elite ABC reagent (Vector). Sections were washed three times in TI buffer (0.05 M Tris pH 7.4, 0.05 M Imidazole (Sigma) in PBS). Signals were amplified using the Tyramide Signal Amplification kit (Perkin Elmer) according to the manufacturer’s recommendations. The sections were then mounted on glass slides and processed through 70%, 90% and 100% alcohols and xylene before cover-slipping with DPX mountant (Sigma). Histological images were obtained using a Zeiss AxioSkop2 plus microscope fitted with a Zeiss AxioCam HRc colour camera. Images were recorded using Zeiss AxioVision 4.7.

### Recruitment assay with fluorescent proteins and immunofluorescence

Free-floating brain sections were prepared as above. Sections were permeabilized in 0.3% PBST and washed three times in PBS. Sections were treated with TrueBlack (Cat No. 23007, Biotium) according to the manufacturer’s recommendations to quench autofluorescence. Control sections were always included to check for complete autofluorescence quenching. Sections were then incubated with exon1 HTT peptides at 50 nM in PBS overnight at 4°C. Rigorous washing in 12 or 24 well plates four to six times of an excess of PBS was performed for six to eight hours at room temperature or for 24-48 hours at 4°C. For immunofluorescence, sections were incubated with primary antibodies (Supplementary Table 1, online resource 1) overnight at 4°C. Sections were washed and incubated with secondary antibodies (all at 1:1000, Thermo Fisher Scientific) for 1 h at room temperature or overnight at 4°C. Sections were counterstained with Hoechst and mounted for confocal microscopy.

### Confocal microscopy and image analysis

Confocal imaging and analysis were performed as previously described with modifications [82]. Images were recorded on a Zeiss LSM710 confocal microscope. Imaging settings were kept consistent for wild-type and HD sections in each experiment. Sample autofluorescence levels were routinely checked by using additional imaging channels. Images were analysed with Fiji (RRID: SCR_002285) and Icy (RRID: SCR_010587) software. The brightness and contrast were adjusted only for presentation purposes.

### Correlative light and electron microscopy (CLEM)

CLEM was performed as described in [32,1] with modifications. Mice were perfused with 4% paraformaldehyde (PFA), brains removed and immediately sectioned into 100 μm slices on a vibratome (Leica Microsystems), then incubated with fluorescently labelled HTT-exon1-43Q in 1 M phosphate buffer at 4 °C. Tissue sections were high-pressure frozen in 100 μm-deep wells of aluminium carriers in the presence of 20% bovine serum albumin (BSA, Sigma) using a HPM100 (Leica Microsystems). Freeze substitution and Lowicryl HM20 (Polysciences, Inc.) resin embedding were performed as previously described [56], and 0.008% uranyl acetate in acetone was used for freeze substitution. Sections of 70-300 nm thickness were cut with an Ultra 35° diamond knife (Diatome) on an ultramicrotome (Leica Microsystems). The sections were picked up with 200 mesh gold NH2 finder grids coated with holey QUANTIFOIL carbon film (hole size = 3.5 μm separated by 1 μm). For high-accuracy CLEM, 50 nm fluorescent TetraSpeck beads (custom order, Thermo Fisher Scientific) were adsorbed onto the grids for fluorescence light microscopy as described [1]. For electron tomography, 10 nm gold particles (Electron Microscopy Sciences) were adsorbed onto both sides of the sections. A Tecnai F20 transmission electron microscope (TEM) (equipped with a Direct Electron DE-20 detector operated at 200 kV) or FEI Polara F30 TEM (fitted with a Quantum energy filter (Gatan) operated in zero-loss imaging mode with a 20 eV energy-selecting slit at 300 kV and a K2 Summit direct electron detector) were used to generate low magnification whole grid maps and intermediate magnification montage maps by Navigator using SerialEM [68,48]. QUANTIFOIL carbon film holes and sample landmarks recognizable in both LM and EM were used for the initial correlation. Further fiducial-based correlation was done using the ec-CLEM plugin in Icy as described [55].

### Immunogold labelling

Immunogold labelling was performed as described in [22] with modifications. Ultrathin sections were prepared on QUANTIFOIL finder grids and fluorescence recorded as described above. Grids were incubated in 0.05 M glycine in Tris-buffered saline (TBS, pH 7.6) for 20 min and then incubated in blocking buffer (5% BSA and 0.1% cold water fish skin gelatine in TBS) for 15 min. After two 5 min washes in blocking buffer, the grids were transferred to incubation buffer (1% BSA and 0.1% cold water fish skin gelatine in TBS) containing anti-HTT antibodies (S830 or mEM48, both at 1:20) and incubated for 48-72 h at 4°C in a humidified chamber. After three washes in the incubation buffer, the grids were incubated with secondary antibodies conjugated to 10 nm or 20 nm gold (ab39609 or ab27242, Abcam, at 1:50 dilution) for 1 h at room temperature. Grids washed three times in TBS and post-fixed in TBS containing 2% glutaraldehyde (Sigma) for 5 min. The grids were rinsed three times in distilled water and post-stained for TEM.

### Electron tomography data collection and processing

Electron tomograms were collected using a FEI Tecnai T12 as previously described [56] or a FEI Tecnai G2 Polara operating at 300 kV and equipped with Quantum energy filter (Gatan) operated in zero-loss imaging mode with a 20 eV energy-selecting slit. Data were recorded on a K2 Summit direct electron detector (Gatan) operating in counting mode at dose rate 5.974 e^-^ per pixel per second with a final sampling of 4.41 Å per pixel. Defocus ranged between −3.5 μm and −6 μm. The dose-symmetric tilt scheme [19] implemented in SerialEM [48] was used to collect tilt series between −60° and 60°, with 3° increments and a total exposure of 600-700 e^-^/Å^2^. 60 frames were collected per tilt and aligned with MotionCor2 version 1.0.5 [81]. Tomograms were reconstructed from tilt series by weighted back-projection using the IMOD package, version 4.9 [31]. Tomography data will be deposited in the Electron Microscopy Data Bank (EMDB).

### Western blotting

Western blotting was performed as previously described [35]. Briefly, mouse hippocampal samples were homogenized in ice-cold IGEPAL RIPA buffer with protease inhibitors at 4 °C. 20 or 40 μg of total protein in Laemmli loading buffer was denatured at 85 °C for 10 min and separated by 7.5-, 10-, or 12% Criterion TGX Stain-free SDS-PAGE (Bio-Rad) and blotted onto 0.45 μm nitrocellulose membrane (Bio-Rad) by submerged transfer apparatus (Bio-Rad). Membranes were then blocked and incubated overnight with gentle agitation at 4 °C with the primary antibody (Supplementary Table 1, online resource 1). Protein was detected by chemiluminescence (Clarity, Bio-Rad) according to the manufacturer’s instructions and signals were captured on a ChemiDoc Touch MP Imaging System (Bio-Rad).

### Quantification and statistical analysis

Statistical analyses were performed by Student’s *t*-test, one-way or two-way ANOVA with Šidák *post-hoc* correction in GraphPad Prism v8. p < 0.05 considered statistically significant; n represents animals or biological replicates. Where shown, ns = not significant, *p < 0.05, **p < 0.01, ***p < 0.001. For all graphs, the bars show mean values, while the error bars represent the standard deviation (SD).

### Database depositions

Representative tomograms have been deposited in the Electron Microscopy Data Bank (EMDB), accession codes are EMD-12374, EMD-12376, EMD-12536 and EMD-12539.

## Results

### Mutant exon 1 HTT proteins are recruited to a subset of the cytoplasmic inclusions in HD mouse brains

It had been demonstrated previously that synthetic, biotinylated polyQ peptides were recruited to aggregation foci in tissue sections from HD *post-mortem* brain and HD mouse models, but not to brain sections from control subjects or wild-type mice [52]. To investigate this phenomenon further, we began by testing whether one of these peptides, 30 glutamines conjugated to biotin via a polyethylene glycol linker (mbPEGQ30), might be recruited to aggregates in brain sections from the zQ175 knockin mouse model of HD. We detected aggregates capable of polyQ peptide recruitment throughout the brain of heterozygous zQ175 mice (Fig. 1a). These recruitment foci did not necessarily co-localize with the HTT inclusions detected by anti-HTT antibodies (Fig. 1b), consistent with the previous report [52]. The S830 anti-HTT antibody identified HTT aggregates in the form of small inclusion bodies in the nucleus and cytoplasm (Fig. 1b, blue arrowheads) as well as a diffuse granular stain that filled neuronal nuclei (Fig. 1b, purple arrows). In contrast, the mbPEGQ30 peptide was recruited to structures that appeared as defined puncta (Fig. 1b, black arrows), and these recruitment foci could be detected in the cortex and hippocampal CA1 region as early as three months of age.

**Fig. 1.**
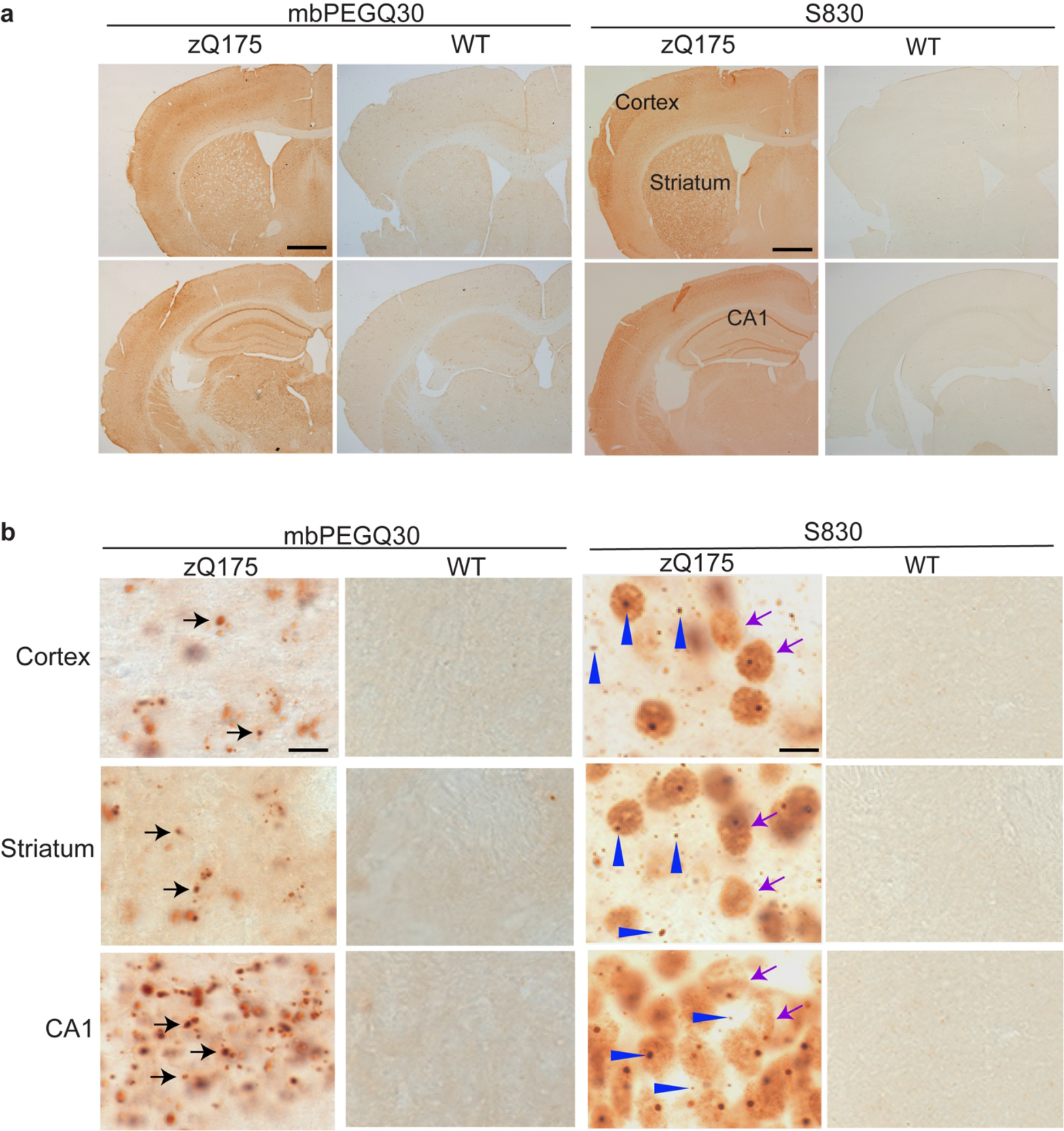
Comparison of mbPEGQ30 peptide recruitment and S830 antibody staining on brain sections from zQ175 mice and their wild-type littermates. **a** Coronal-hemispheres from 5-month-old zQ175 mice and their wild-type littermates, either treated with the mbPEGQ30 peptide or immunolabelled with the S830 antibody (2.5x). **b** Zoomed-in view of cortical, striatal and hippocampal (CA1) regions from 5-month-old zQ175 and wild-type mice, either treated with the mbPEGQ30 peptide or immunolabelled with S830. Black arrows in **b** point to the defined puncta of recruitment signal. Purple arrows indicate the diffuse nuclear aggregation; blue arrowheads point to small inclusion bodies. WT = wild-type. Scale bars: **a** 400 μm; **b** 10 μm.

To investigate the nature of the recruitment foci, we set out to determine whether these structures could recruit a fluorescently labelled exon 1 HTT protein. Exon 1 HTT, with a polyQ stretch of 43 glutamines (HTT-exon1-43Q), was synthesized as previously described [73,76], and conjugated at the C-terminus with either an Alexa Fluor 488 (AF488) or Alexa Fluor 647 (AF647) dye. The fluorescently labelled exon 1 HTT proteins were applied to brain sections from zQ175 mice at 6 months of age and from R6/2 mice at 12 weeks of age, together with their respective wild-type littermate controls. At these ages, the zQ175 mice are presymptomatic and yet to present overt behavioral phenotypes [49,20], with end-stage disease not occurring until approximately 20 months of age. However, R6/2 mice are symptomatic, having developed pronounced motor and cognitive impairments before 8 weeks of age [59,43]. HTT-exon1-43Q was recruited to structures distributed throughout the brain sections for both zQ175 and R6/2 mice (Fig. 2a, b). The fluorescent signal was not quenched by either pre- or post-treatment with Trueblack (Supplementary Fig. 1, online resource 1), indicating that it did not represent autofluorescence. Next, we immunoprobed HTT-exon1-43Q treated sections with the S830 anti-HTT antibody and examined them for evidence of colocalization in the cortex, striatum and hippocampus. S830 detects nuclear aggregation in the form of a diffuse granular pattern as well as nuclear inclusions; it also labels cytoplasmic aggregates in the form of inclusion bodies that are perinuclear or scattered throughout the neuropil [9,39] (Fig. 2a, b). Colocalization of HTT-exon1-43Q with S830 fluorescent signals was confined to perinuclear inclusions in both zQ175 and R6/2 mice (Fig. 2c).

**Fig. 2.**
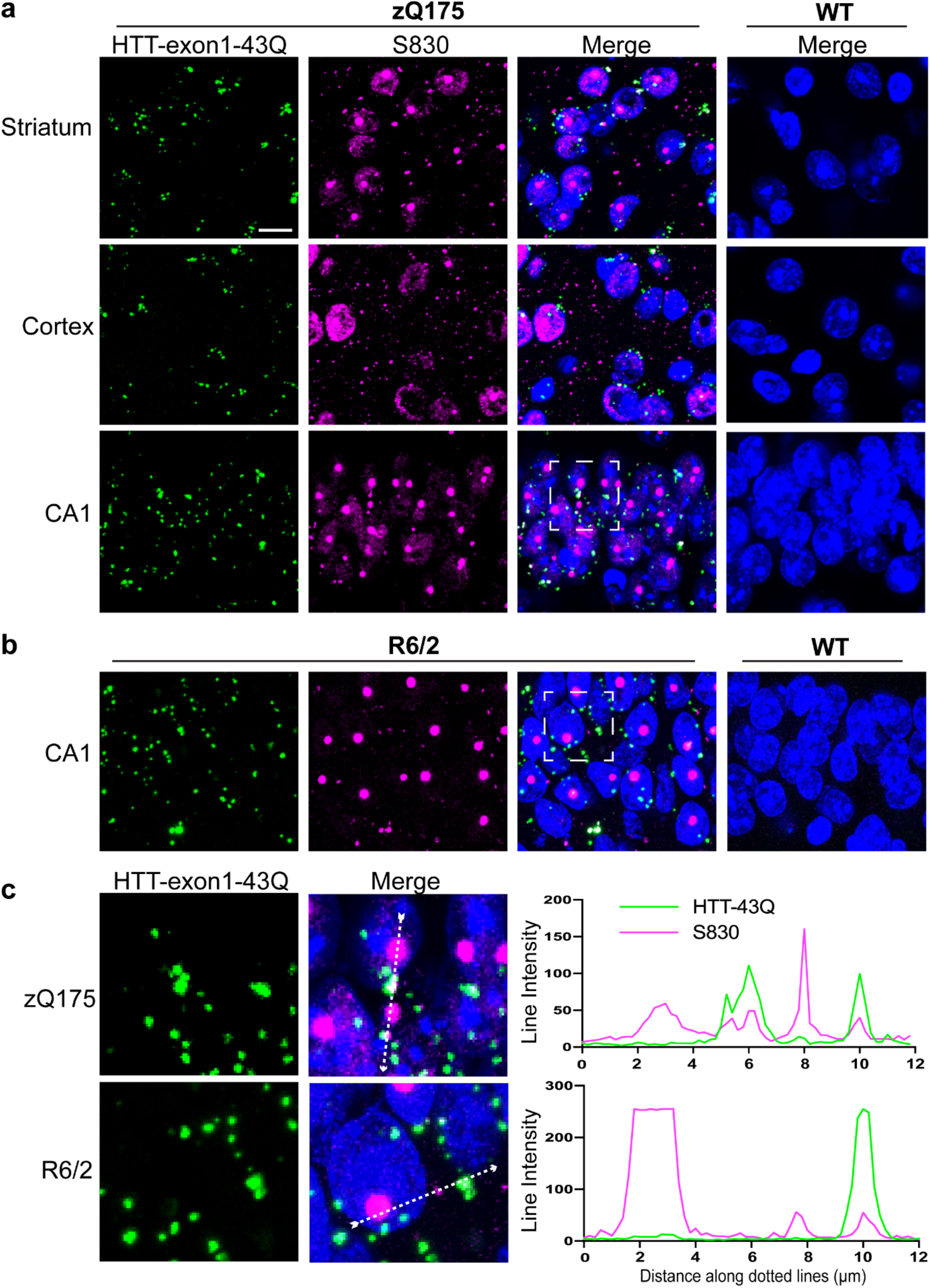
Co-localization of HTT-exon1-43Q signal with S830 antibody immunostaining**. a** HTT-exon1-43Q (green) was applied to TrueBlack treated zQ175 and wild-type brain sections from 6-month-old mice which were immunoprobed with the S830 antibody (magenta) and nuclei were counterstained with Hoechst (blue). HTT-exon1-43Q was recruited to perinuclear inclusions in the striatum, cortex, and hippocampus (CA1). **b** R6/2 and wild-type brain sections from 12-week-old mice treated with the HTT-exon1-43Q protein were immunoprobed with S830 and nuclei were counterstained with Hoechst. HTT-exon1-43Q was recruited to perinuclear inclusions as shown in the hippocampal CA1 region. **c** Enlarged areas from white boxed regions in **a** and **b**, respectively. The intensity profiles were plotted along the white dotted lines. The HTT-exon-43Q protein was not recruited to nuclear inclusions. Scale bars: 10 μm. WT = wild type.

### Recruitment of the exon 1 HTT proteins to HD mouse brain sections was both disease-stage and polyQ-length dependent

To determine whether the extent of HTT-exon1-43Q recruitment to brain sections was dependent on disease progression, we applied HTT-exon1-43Q to coronal sections from R6/2 mice and their wild-type counterparts at 4, 8 and 12 weeks of age. The recruitment signal was prominent on sections from both 8- and 12-week-old mice (Fig. 3a). Although barely detectable by eye, (Fig. 3a), quantification revealed a significant increase in the fluorescence intensity of HTT-exon1-43Q treated sections from R6/2 as compared to wild-type mice at 4 weeks of age (Fig. 3c). The recruitment foci intensities significantly increased in R6/2 mice with disease progression (Fig. 3c) and HTT-exon1-43Q was not recruited to sections from wild-type mice at any of these ages (Fig. 3a, c).

**Fig. 3.**
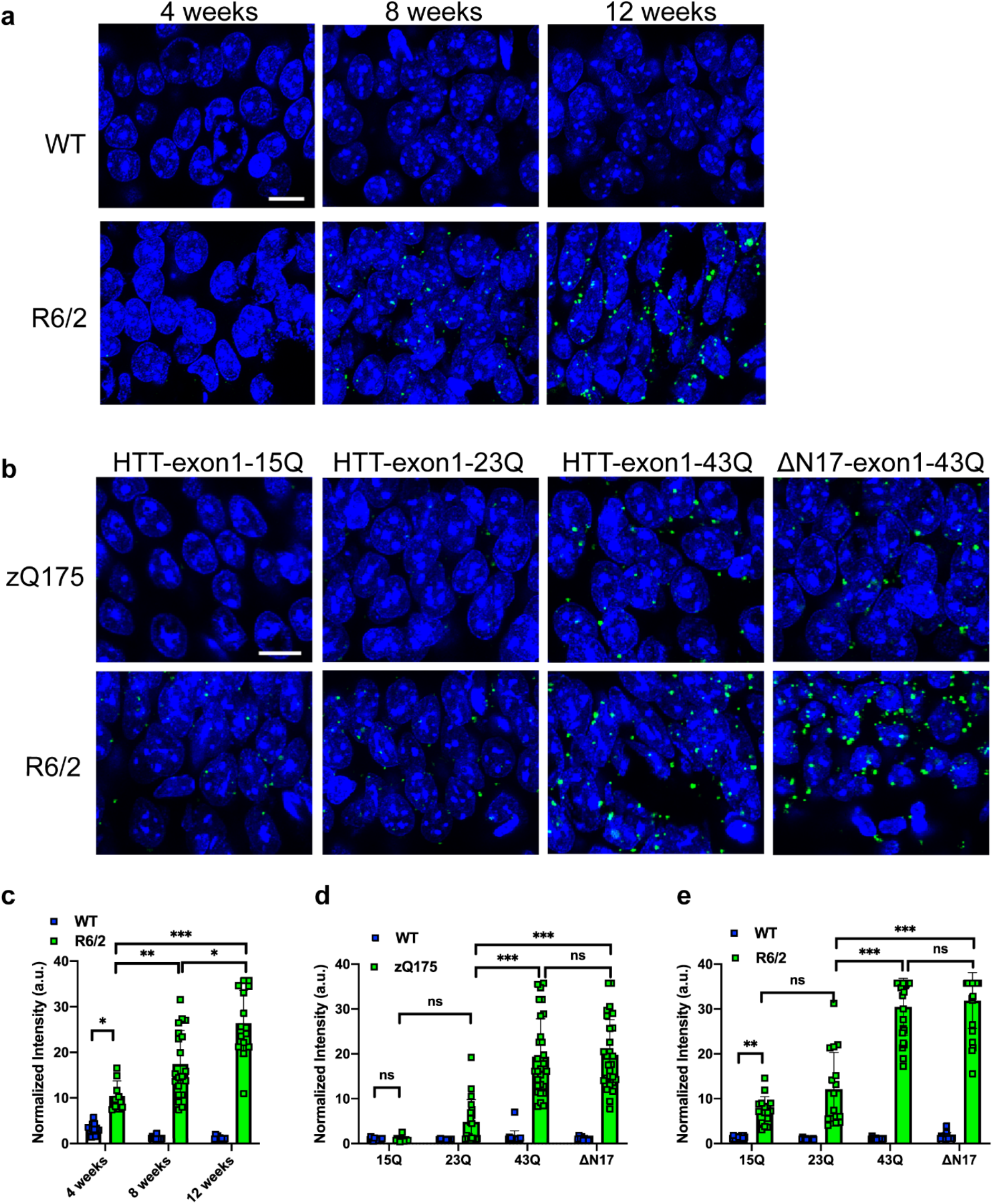
Recruitment of exon 1 HTT proteins to brain sections from HD mouse models is disease-stage and polyQ-length dependent. **a** Representative confocal microscopy images for recruitment of HTT-exon1-43Q to the hippocampus (CA1) of R6/2 and wild-type mice at 4, 8 and 12 weeks of age. Nuclei were counterstained with Hoechst. **b** Recruitment of HTT-exon1-15Q, HTT-exon1-23Q and HTT-exon1-43Q, as well as HTT-exon1-43Q with an N-terminal 17 amino acid truncation (ΔN17-exon1-43Q) to 6-month-old zQ175, and 12-week-old R6/2, brain sections. Nuclei were counterstained with Hoechst. **c** Quantification of fluorescence intensities of HTT-exon1-43Q treatment for R6/2 and wild-type mice at 4, 8 and 12 weeks of age. **d, e** Quantifications of fluorescence intensities of **d** zQ175 and **e** R6/2 sections treated with exon 1 HTT proteins with different polyQ length. Statistical analysis was by two-way ANOVA. Data represented by mean ± S.D., two sets of 15-25 images for three replicates per experiment. *p<0.05; **p < 0.01; ***p < 0.001, ns = not significant. Scale bars: 20 μm. WT = wild-type.

To investigate which structural domains within exon 1 might be important for the HTT-exon1-43Q recruitment, we generated exon 1 HTT monomers with a range of polyQ lengths (15Q, 23Q, 43Q) and the HTT-exon1-43Q peptide lacking the first 17 amino acids (ΔN17-exon1-43Q). The first 17 amino acids of HTT are known to oligomerize through α-helix-rich interactions [71,26] and the polyQ tracts interact though β-strand structures [21]. These proteins were applied to coronal sections from zQ175 mice at 6 months and R6/2 mice at 12 weeks of age, respectively. Treatment of the zQ175 brain sections with HTT-exon1-15Q gave a comparable signal to the wild-type controls (Fig. 3b, d), and therefore 15 glutamines were not sufficient for recruitment on 6-month-old zQ175 brain sections. Whilst some AF488 signal was obtained with the HTT-exon 1-23Q peptide on zQ175 sections, the fluorescence intensities were not significantly increased over HTT-exon1-15Q (Fig. 3b, d). Treatment of the zQ175 brain sections with either HTT-exon1-43Q or ΔN17-exon1-43Q resulted in significantly increased fluorescence intensities compared to HTT-exon1-15Q and HTT-exon1-23Q (Fig. 3b, d). In contrast to 6-month-old zQ175 mice, both HTT-exon1-15Q and HTT-exon1-23Q were recruited to R6/2 sections from 12-week-old mice (Fig. 3b, e). The extent of recruitment was comparable for the HTT-exon1-43Q and ΔN17-43Q proteins (Fig. 3b) on both zQ175 (Fig. 3d) and R6/2 (Fig. 3e) sections. Therefore, the recruitment of exon 1 HTT proteins increased with disease progression and aggregate burden, and this interaction occurred through the polyQ tract.

### HTT-exon1-43Q proteins are recruited to single membrane bound, vesicle-rich organelles

We next established a protocol for correlative light and electron microscopy (CLEM) to determine the ultrastructure of the recruitment foci. Coronal sections were prepared from zQ175 mice at 6 months of age, immediately after perfusion. The HTT-exon1-43Q-AF647 peptide was applied to the zQ175 sections, which were then subjected to high-pressure freezing and freeze substitution, and the recruitment sites in the hippocampal CA1 region were imaged. We found that the HTT-exon1-43Q-AF647 fluorescence signal was well-preserved following these procedures (Fig. 4a, c, magenta), and the autofluorescence of the CA1 region was negligible as shown by auto-exposure of the sample in the 488 nm channel (Fig. 4b). Ultrathin sections were collected on QUANTIFOIL finder EM grids and fluorescence light microscopy (LM) images were recorded (Fig. 4e). As shown in Fig. 4g, the recruitment sites were identified as single membrane bound, vesicle rich organelles (see Supplementary Fig. 2, online resource 1 for larger view of the organelle and Supplementary Video 1, online resource 2 for tomogram). In this example, the recruitment organelle (magenta) was adjacent to a double membrane, phagophore-like structure, as shown in the 3D segmentation from the tomogram (Fig. 4h, cyan).

**Fig. 4.**
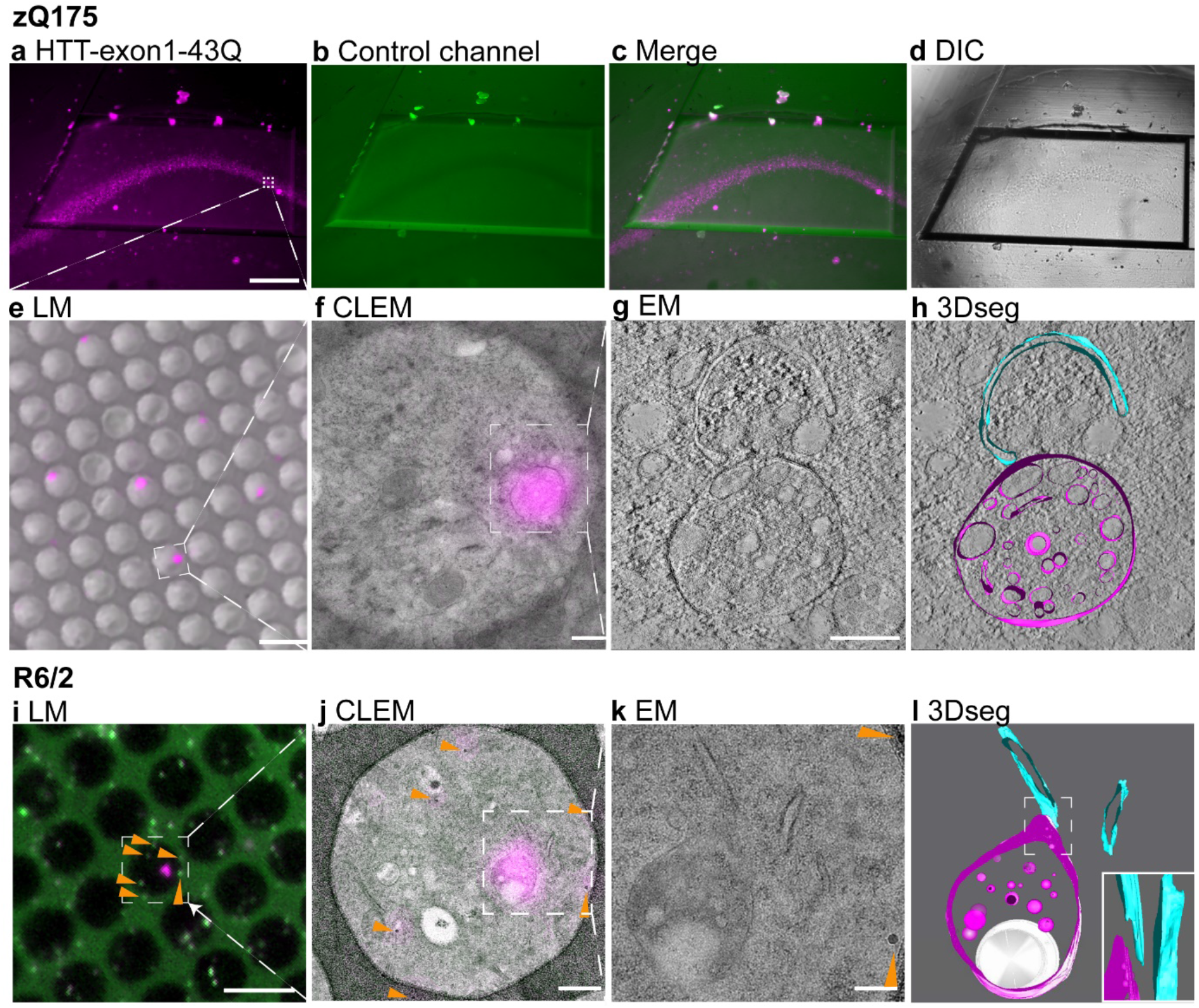
CLEM reveals that the exon 1 HTT proteins are recruited to single membrane bound organelles. **a-d** 6-month-old zQ175 coronal sections were treated with the HTT-exon1-43Q-AF647 protein and subjected to high pressure freezing and freeze substitution. **a-c** Fluorescence images were taken at 10x magnification using settings for **a** Alexa Fluor 647 showing the HTT-exon1-43Q-AF647 signal in magenta and **b** Alexa Fluor 488 to show autofluorescence levels. **c** merged image from **a** and **b**. **d** DIC at 10x magnification. **e-h** Ultrathin sections were subsequently prepared and placed on holey carbon QUANTIFOIL EM finder grids. **e** LM image including both fluorescence and DIC. **f** white boxed area in **e** showing the overlay of the fluorescent signal with EM. **g** 2D EM image from the white box in **f**. **h** the corresponding tomogram section of **g** with segmentation of the tomogram shown in Supplementary Video 1, online resource 2. The recruitment signal (magenta) labelled a single membrane bound, vesicle rich organelle, next to a double membrane, phagophore-like structure (cyan). **i-k** CLEM of HTT-exon1-43Q on 12-week-old R6/2 sections with TetraSpeck beads. **i** LM image of an ultrathin brain section on a QUANTIFOIL grid with 50 nm TetraSpeck beads (shown as green puncta and indicated by orange arrowheads). **j** White-boxed area in **i** showing the overlay of fluorescent signal with EM. Orange arrowheads indicate the TetraSpeck beads. **k** EM image of the structure in the white boxed area in **j**. **l** the tomogram of the structure in **k** was used to generate this 3D segmentation. The recruitment signal (magenta) uncovered a single membrane-bound vesicle-rich organelle containing a large vacuole. This was adjacent to a double membrane structure (cyan). Solid white boxed insert is a side view of the dashed white box area. Scale bars: **a** = 100 μm, **e** = 5 μm, **f** = 300 nm, **g** = 200 nm, **i** = 5 μm, **j** = 450 nm, **k** = 200 nm. DIC = differential interference contrast, LM = light microscopy, CLEM = correlative light and electron microscopy, EM = electron microscopy, 3Dseg = 3D segmentation.

We used fluorescent fiducials to provide markers in the field of view to accurately map fluorescence to electron micrographs, achieving an accuracy of less than 100 nm [32,1]. TetraSpeck beads have a wide fluorescence spectrum and can be identified in both fluorescence channels (Fig. 4i, green and magenta) [1]. The same sets of beads could be observed by both LM and EM as indicated by orange arrows in Fig. 4i-k. Whereas the beads were visible in multiple channels by LM, the recruitment signal was only seen in one channel (Fig. 4i), so that the region of interest (ROI) (magenta) could be readily identified.

Tomography and 3D segmentation revealed that on this R6/2 section, HTT-exon1-43Q was again recruited to a single membrane bound, vesicle-rich structure containing a large vacuole (Fig. 4k-l). The organelle was close to a double membrane structure, either part of the endoplasmic reticulum (ER) or, once again, phagophore-like. Tomography and segmentation established that the organelle was in close proximity to, but not in direct contact with, this double membrane structure (bottom right insert in Fig. 4l). These CLEM results demonstrated that the HTT-exon1-43Q proteins were recruited to similar, single membrane bound, vesicle-rich organelles in both zQ175 and R6/2 mice.

### Mutant HTT is localized to the lumen of the recruitment organelles

In order to determine which structures within the recruitment organelles contain mutant HTT, we used immunogold labelling on R6/2 brain sections from 12-week-old mice. Two anti-HTT antibodies were used: S830, labelled with 10 nm, and mEM48 labelled with 20 nm gold particles, respectively. Both S830 (Fig. 5a, c-f) and mEM48 (Fig. 5b) labelled mutant HTT within the single membrane-bound organelles. In both cases, the gold particles were located in the lumen, especially in the electron lucent regions (Fig. 5d, f), but were absent from the large vacuoles (Fig. 5b, e). Three independent experiments revealed that, on average, the lumen of 93% of organelles examined contained S830 gold particles (in total 45 out of 47 ROI) (Fig. 5g). The mEM48 labelling was sparser, with 61% of the organelles containing gold particles (Fig. 5g). The immunogold staining background was relatively clean for both S830 and mEM48, with very few gold particles identified in the non-ROI area of the same QUANTIFOIL carbon hole (Fig. 5h). Although fibrillar material is well preserved in high pressure frozen, freeze-substituted samples, no fibrils were observed in the recruitment organelles.

**Fig. 5.**
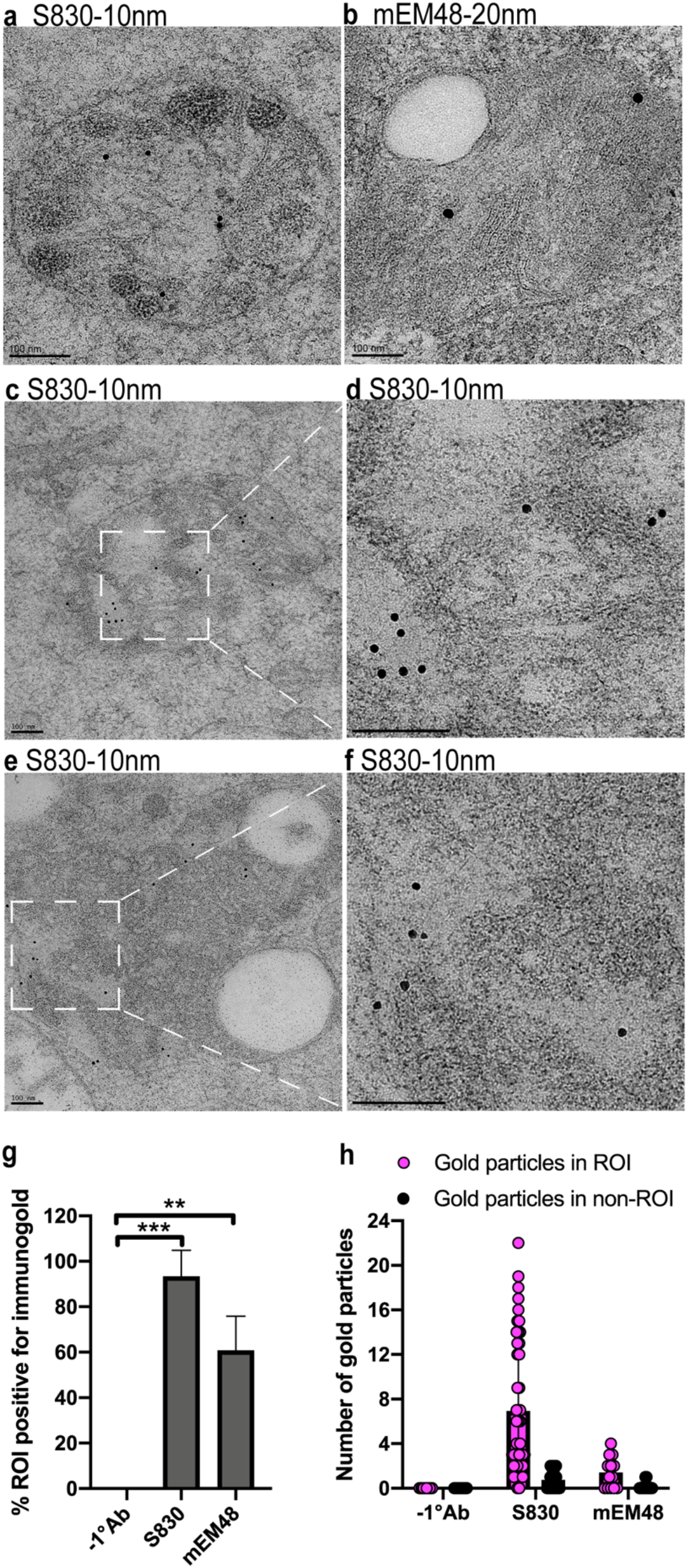
Mutant HTT is localized to the electron-lucent areas within the lumen of the organelles. **a-f** Immunogold-labelled S830 and mEM48 antibodies were imaged on EM sections from 12-week-old R6/2 mice **a, c-f** shows organelles labelled with S830 10 nm gold and **b** with EM48 20 nm gold. **d, f** Enlarged views of the boxed areas in **c** and **e**, respectively, showing that the gold particles are localized to electron-lucent areas inside the organelles. **g** Percentage of organelles that contained gold particles after immunogold labelling. Three independent experiments were performed for S830 and two for mEM48, with approximately 15 micrographs being imaged for each experiment. **h** Relative numbers of gold particles inside the organelle ROI compared to those in non-ROI areas of the same QUANTIFOIL carbon hole. Statistical analysis was by Student’s *t*-test. Data represented by mean ± S.D. **p < 0.01; ***p < 0.001. Scale bars: 100 nm. ROI = region of interest. −1°Ab = without primary antibody.

### The single-membrane organelles differ between presymptomatic and late-stage HD

Our CLEM investigations revealed similar recruitment organelle structures for the zQ175 knockin and R6/2 transgenic mouse models of HD. In both cases, they were single membrane-bound organelles, often containing vesicles. Those with many intraluminal vesicles (ILVs) resembled multivesicular bodies (MVB) or amphisomes (Fig. 6a, b, e and Supplementary Video 2, online resource 3) [14,8,10,79,64]. Others, with prominent electron lucent vacuole(s), sometimes contained internal membranous structures (luminal lamellae), and resembled lysosomal organelles such as autolysosomes or residual bodies (Fig. 6c, d, f-h, Supplementary. Video 3, online resource 4 and Supplementary Video 4, online resource 5) [38,6,33,30]. Detailed analysis revealed that in zQ175 mice, on average, 71% of the recruitment organelles were MVBs/amphisomes while 29% were autolysosomes/residual bodies. In R6/2 mice, the number of MVBs/amphisomes was on average 38%, and the number of autolysosome/residual bodies was 62% (Fig. 6i).

**Fig. 6.**
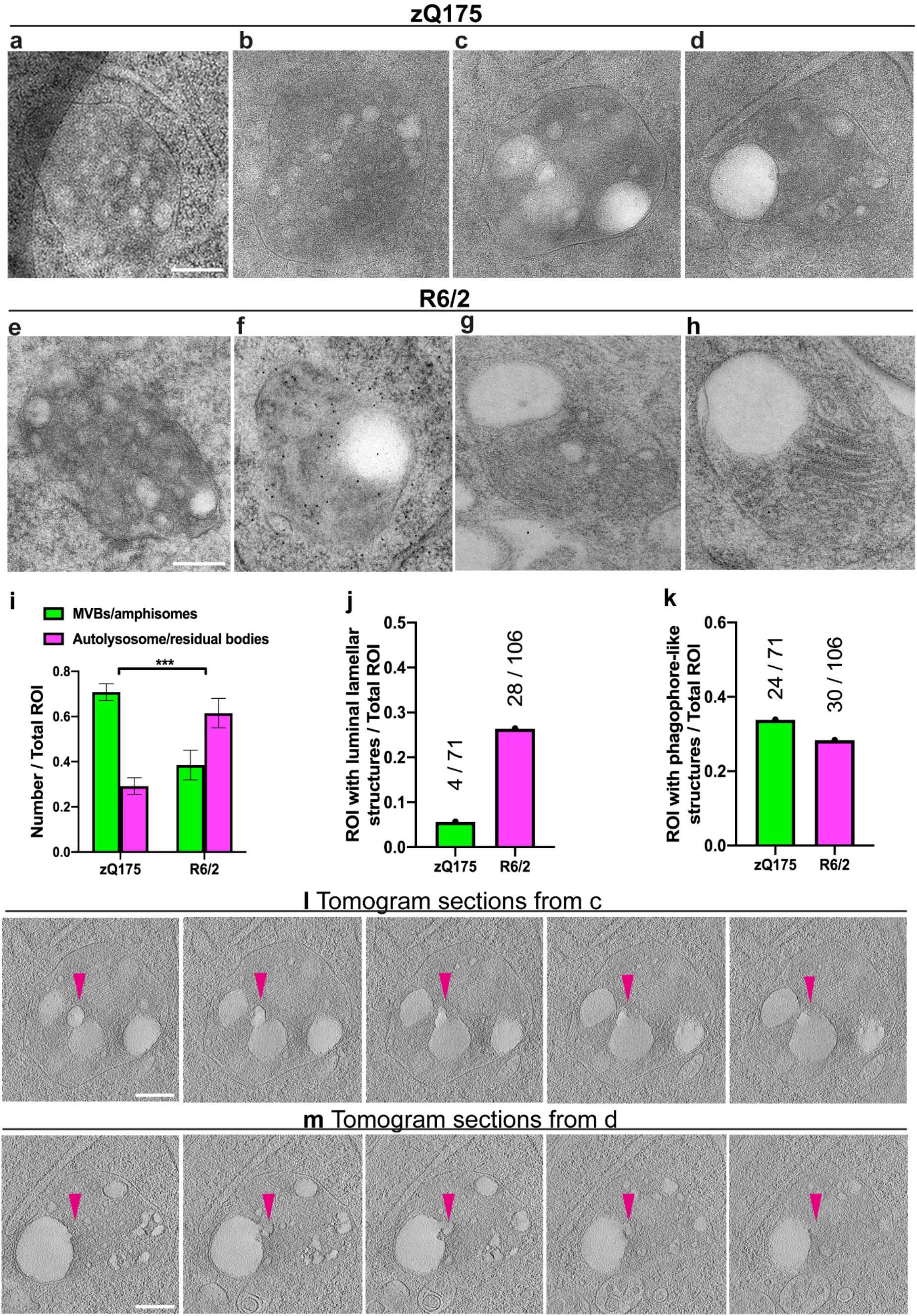
Ultrastructural analysis in zQ175 and R6/2 mice. **a-h** Representative recruitment foci in 6-month-old zQ175 and 12-week-old R6/2 mice. **i** Quantification of the number of MVB and autolysosome/residual bodies in zQ175 and R6/2 sections. **j** Quantification of the ratio between ROI with intraluminal lamellar structures and the total ROI. **k** Quantification of ROI with phagophore-like double membrane structures as a ratio of the total number of ROI. Four independent experiments with a total of 71 and 106 foci examined on TEM images for zQ175 and R6/2, respectively. Statistical analysis was two-way ANOVA, data represented by mean ± S.D. ***p < 0.001. **l**, **m** Serially displayed sections of two tomograms from **c** and **d**, presented as sequences taken as every 10th slice through the 8x binned reconstructed tomograms. Related tomograms are shown in Supplementary Video 3, online resource 4 and Supplementary. Video 4, online resource 5. Scale bars: 200 nm.

Luminal lamellae were observed in some recruitment organelles in both zQ175 (Fig. 6c and Supplementary Video 3, online resource 4) and R6/2 (Fig. 6g, h) mice. In zQ175, only 6% of organelles (4 out of 71) had lamellar structures, whereas in R6/2, this number was 26% (28 out of 106) (Fig. 6j). Double membrane, phagophore-like structures were frequently observed in close proximity to the recruitment organelles. In zQ175 mice, 34% (24 out of 71) of the organelles had an adjacent double membrane structure and in R6/2 mice, this was 28% (30 out of 106) (Fig. 6k).

Sequential sections through the 3D volumes of reconstructed tomograms suggest that vesicles fuse to form larger vacuoles within the recruitment organelles (Fig. 6l, m, Supplementary. Video 3, online resource 4 and Supplementary Video 4, online resource 5).

### The recruitment organelles are members of the endolysosomal pathway

We next confirmed by confocal fluorescence microscopy that the recruitment organelles identified by CLEM were a subset of endolysosomal organelles. HTT-exon1-43Q proteins were applied to brain sections from 6-month-old zQ175 mice, which were then immunoprobed for the lysosomal-associated membrane protein 1 (LAMP1), a marker which labels MVBs through to lysosomal organelles [7], and the anti-HTT antibody, S830. As indicated by white arrowheads (Fig. 7a-d), and intensity profiles (Fig. 7e, f), the HTT-exon1-43Q recruitment signal colocalized with a subset of puncta that were LAMP1 and S830 positive. LAMP1 was also found to reside in numerous subcellular locations that were negative for the recruitment signal (Fig. 7b, e, f). Therefore, the HTT-exon1-43Q protein was recruited to a subset of LAMP1 positive endolysosomal organelles.

**Fig. 7.**
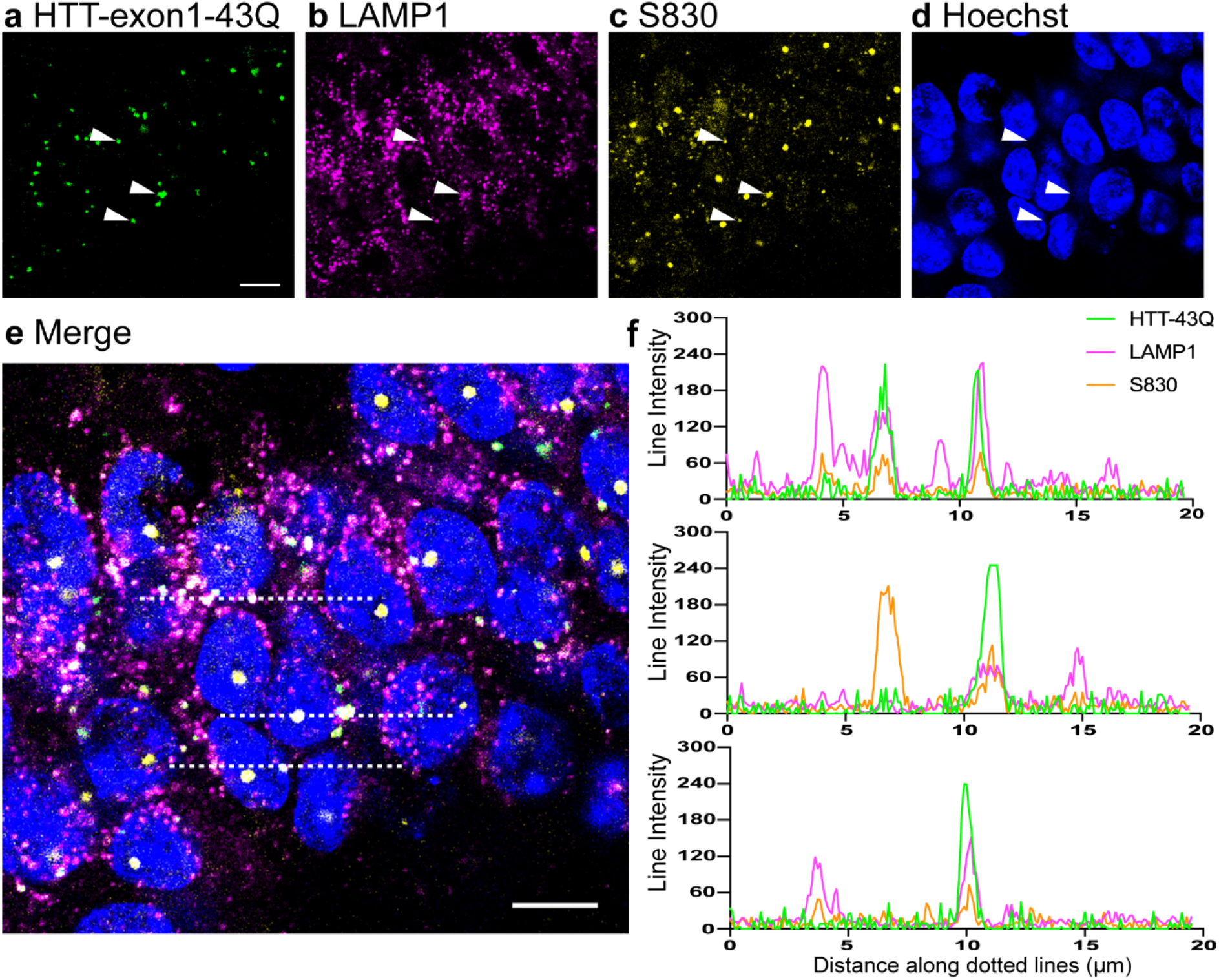
The recruitment organelles are members of a dysregulated endolysosomal system. **a-e** Confocal images of the CA1 region from 6-month-old zQ175 mice showing that HTT-exon1-43Q was recruited to organelles that were also immunostained with an antibody to LAMP1 and with S830. White arrowheads in **a**-**d** indicate the same three regions across different channels. **e** Merged image of all channels from **a**-**d**. The three dotted lines run through the puncta indicated by the arrowheads. **f** Intensity profiles showing the signal along the dotted lines (from left to right) in **e**. Scale bars: 10 μm.

### Dysregulation of endolysosomal system in presymptomatic and late-Stage HD

We next examined whether the endolysosomal system had been altered by measuring total protein levels of endolysosomal markers that recognize early endosomes through to autolysosomes in zQ175 and R6/2 mice and their wide-type littermates. Antibodies to RAB5 (early endosome), RAB7 and M6PR (MVB/late endosome), RAB 11 (recycling endosomes), RAB27 and CD63 (exosomes), LAMP1 (MVBs to autolysosomes) and cathepsin B and cathepsin D (lysosomal proteases) were used to probe western blots of hippocampal lysates from R6/2 mice and their wild-type littermates at 4, 8 and 12 weeks of age (Supplementary Fig. 3, online resource 1). Reductions in LAMP1, RAB7, RAB27 and CD63, and an increase in cathepsin B were identified in R6/2 lysates as compared to wild-type (Supplementary Fig. 3, online resource 1). LAMP1, RAB7, RAB27, CD63 and cathepsin B antibodies were then used to determine whether these proteins were altered in hippocampal lysates from zQ175 mice at 6 months of age, and the western blots of hippocampal lysates from 8- and 12-week-old R6/2 mice were repeated with a larger number of samples. We found that levels of RAB27 and CD63, both involved in MVB related exocytosis [53,23,80], were reduced in the 6-month-old zQ175 and 8- and 12-week-old R6/2 lysates (Fig. 8). Cathepsin B was increased in the 8- and 12-week R6/2 lysates, but not in those from zQ175 mice at 6 months of age (Fig. 8). To conclude, our data suggest that the endolysosomal system is dysregulated in HD and that this occurs at an early, presymptomatic stage.

**Fig. 8.**
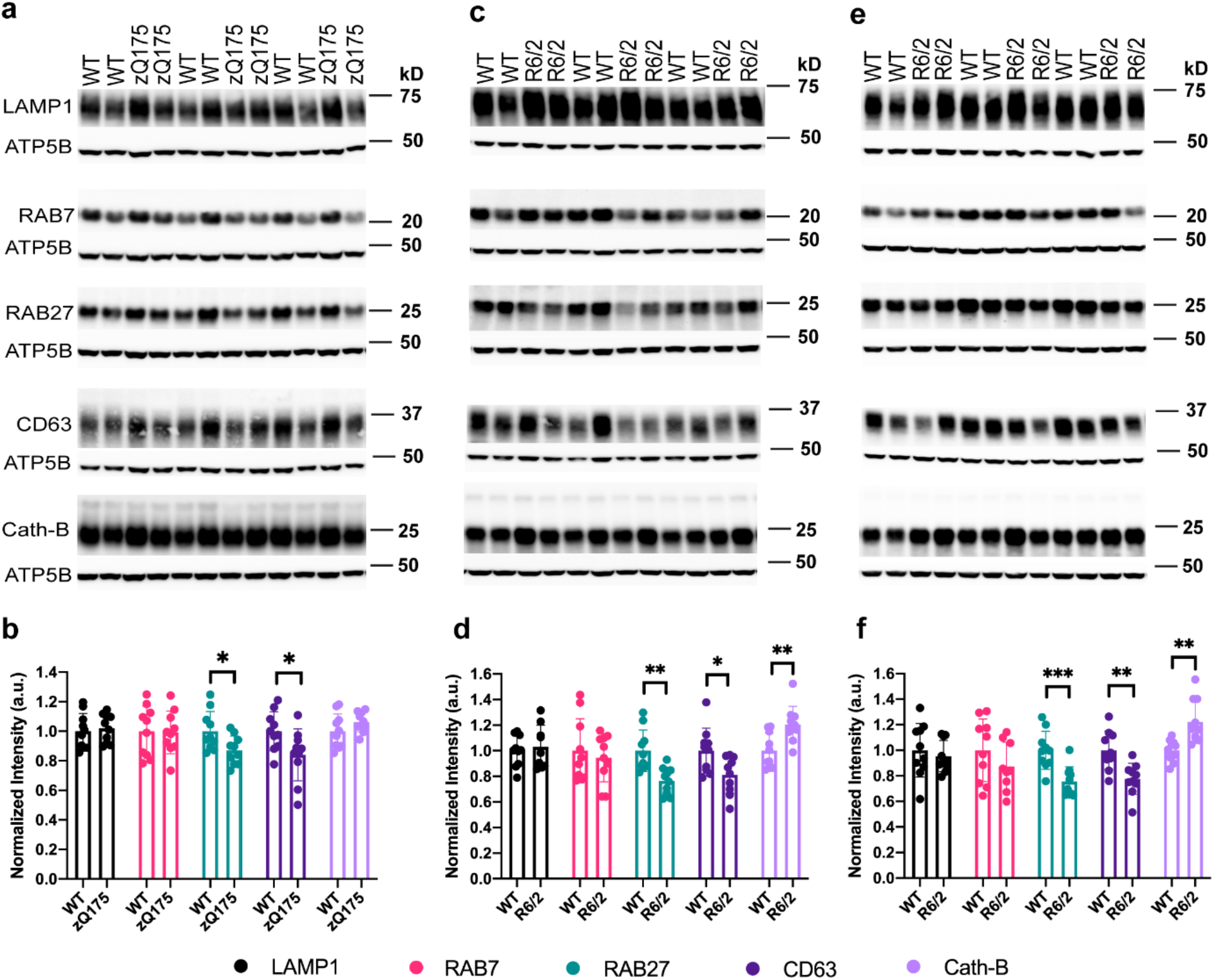
Western blots of endolysosomal proteins in the zQ175 and R6/2 hippocampus. **a** Representative western blots of total protein hippocampal lysates from 6-month-old zQ175 mice and their wild-type littermates immunoprobed for LAMP1, RAB7, RAB27, CD63 or Cathepsin B (Cath-B). ATP5B served as loading control. **b** Quantification of the blot shown in **a**. **c** Western blots of total protein hippocampal lysates from 8-week-old R6/2 mice and their wild-type littermates immunoprobed for the same proteins as in **a**. **d** Quantification of the blot shown in **c**. **e** Western blotting of total protein hippocampal lysates from 12-week-old R6/2 mice and their wild-type littermates immunoprobed for the same proteins as in **a**. **f** Quantification of the blot shown in **e**. Statistical analysis was by Student’s *t*-test. n = 10 / genotype / age. The uncropped blots are shown in Supplementary Fig. 5-7, online resource 1. Data represented by mean ± S.D. *p<0.05; **p < 0.01; ***p < 0.001. Cath-B = Cathepsin B, WT = wild-type.

## Discussion

Precise CLEM targeting of pathological mutant HTT deposits in mouse models of HD has revealed a structure and cellular pathway that were entirely unpredicted by previous *in vitro* and cell models of HTT aggregation. We show that a subset of cytoplasmic antibody-detected ‘inclusions’ in the brains of HD mouse models are single membrane-bound, vesicle-rich organelles. These predominantly resemble MVBs or amphisomes in presymptomatic mice and are more likely to resemble autolysosomes or residual bodies at later stages of the disease course. Immunogold labelling localized mutant HTT to non-fibrillar, electron-lucent regions within the lumen of these organelles. We discovered that proteins important for exosome function were depleted in presymptomatic mice providing further evidence that the endolysosomal system is dysregulated in HD.

In this study, we set out to uncover the structures in tissue sections from mouse models of HD to which polyQ peptides are recruited. To facilitate the potential application of CLEM we tested the use of exon 1 HTT proteins chemically conjugated with fluorescent tags. We utilized two complementary mouse models: R6/2 mice that are transgenic for a human exon 1 *HTT* [47] and the zQ175 knockin HD mouse model [49] with comparable mean CAG repeat expansions of 182 and 192 CAGs, respectively. R6/2 mice progress to end-stage disease by around 14 weeks, whereas disease progression in zQ175 mice is more protracted, with end-stage disease occurring at approximately 20 months of age. We studied the zQ175 mice at 6 months of age prior to symptom onset and analyzed R6/2 mice over their disease course up to 12 weeks, allowing us to study late-stage disease in the absence of confounds due to ageing. Overall, we demonstrated that recruitment of the HTT-exon1-43Q protein occurred through the polyQ tract, in an N17 independent manner, and increased with disease progression. We showed that in all cases, exon 1 HTT proteins with 15Q, 23Q and 43Q polyQ repeat lengths were recruited to sections from 12 week-old R6/2 mice, consistent with the ability of aggregates, once amyloid growth has been nucleated, to bind proteins containing polyQ tracts whether or not the polyQ is in the HD pathogenic range [27].

At the resolution of confocal microscopy, antibody staining identifies inclusion bodies in neuronal nuclei and throughout the neuropil in the brains of HD mouse models and HD patients [9,11,18,39]. On the nanoscale, nuclear inclusions are compact, membrane-less fibril-containing assemblies [9,72]. Consistent with the recruitment of the biotinylated polyQ peptide [52], we found that the HTT-exon1-43Q peptide was not recruited to nuclear inclusions, but instead, to a subset of cytoplasmic sites in the cell body. Structures of polyQ containing aggregates have previously been identified by CLEM in yeast cells, immortalized cell lines and primary mouse neurons that overexpress an expanded polyQ, showing that the deposition of aggregates depends greatly on the cellular environment and context [56,5,17,61]. Application of correlative microscopy to tissue sections is much more challenging. Previously, it has been used to uncover the structure of Lewy bodies in Parkinson’s disease *post-mortem* brain sections, by correlating LM and EM images on adjacent sections, revealing vesicle-rich, mainly non fibrillar aggregates [69]. Unfortunately, this method was not applicable for our study due to the challenge of fluorescent labelling after osmium tetroxide treatment and the smaller size of the recruitment foci. Instead, we successfully adapted a high-precision CLEM methodology originally developed for mammalian and yeast cells [1,32]. This allowed us to correlate LM and EM on the same mouse brain sections at an accuracy of <100 nm, consistent with the cell-based studies [32,1].

Our CLEM analysis revealed that these recruitment foci were single membrane-bound, vesicle-rich structures. Those with many intraluminal vesicles (ILVs) resembled MVBs or amphisomes [14,8,10,79,64] and others, containing vacuole(s) and in some cases internal membranous structures (luminal lamellae), resembled autolysosomes or residual bodies [38,6,33,30]. The presence of luminal lamellae has been associated with various lysosomal storage diseases [10,34]. Here we found more organelles containing luminal lamellae in late-stage HD, possibly due to the accumulation of cellular lipids in lysosomes as a result of lysosomal dysfunction. However, we cannot exclude the possibility that some of the membranous structures came directly from the cellular organelles/debris engulfed by autophagy. Therefore, the exact mechanism of lamellae formation, and how it relates to HD, requires further investigation. The proportions of MVB-like structures and autolysosomes in zQ175 and R6/2 brains at 6 months and 12 weeks of age, respectively, suggested that the MVBs / amphisomes transition to autolysosomes / residual bodies during disease progression. In support of this, examination of membrane contacts in the 3D reconstructions provided evidence of vesicle fusion to form the large vacuole within the autolysosomes. Double membrane phagophore-like structures were frequently observed in close proximity to the recruitment organelles in both zQ175 and R6/2 mice, an association that might be underestimated. TEM images of the sections were used to estimate the number of cases in which phagophore-like double membranes were associated with the organelles. Since some associated phagophore-like structures might lie outside the plane of the section, the true number of associations could be greater. On the other hand, we cannot exclude the possibility that some of these double membranes are smooth ER. Future studies with large volume 3D EM reconstruction will be valuable in further clarifying this association.

The endolysosomal pathway links exosome biogenesis and autophagy and is important in responding to stress and maintaining homeostasis in health and disease [79,13] (Fig. 9). Early endosomes mature into MVBs which are late endocytic compartments containing many ILVs. MVBs can fuse with the plasma membrane, releasing the intraluminal vesicles into the extracellular space as exosomes. Through the process of macroautophagy, cytoplasmic proteins and organelles are engulfed by the double membraned phagophore to form autophagosomes, which fuse with lysosomes to degrade the contents. Alternatively, autophagosomes can fuse with MVBs to form hybrid organelles termed amphisomes, which are thought to eventually fuse with lysosomes. Amphisomes can also fuse with the plasma membrane and secrete their contents (Fig. 9a).

**Fig. 9.**
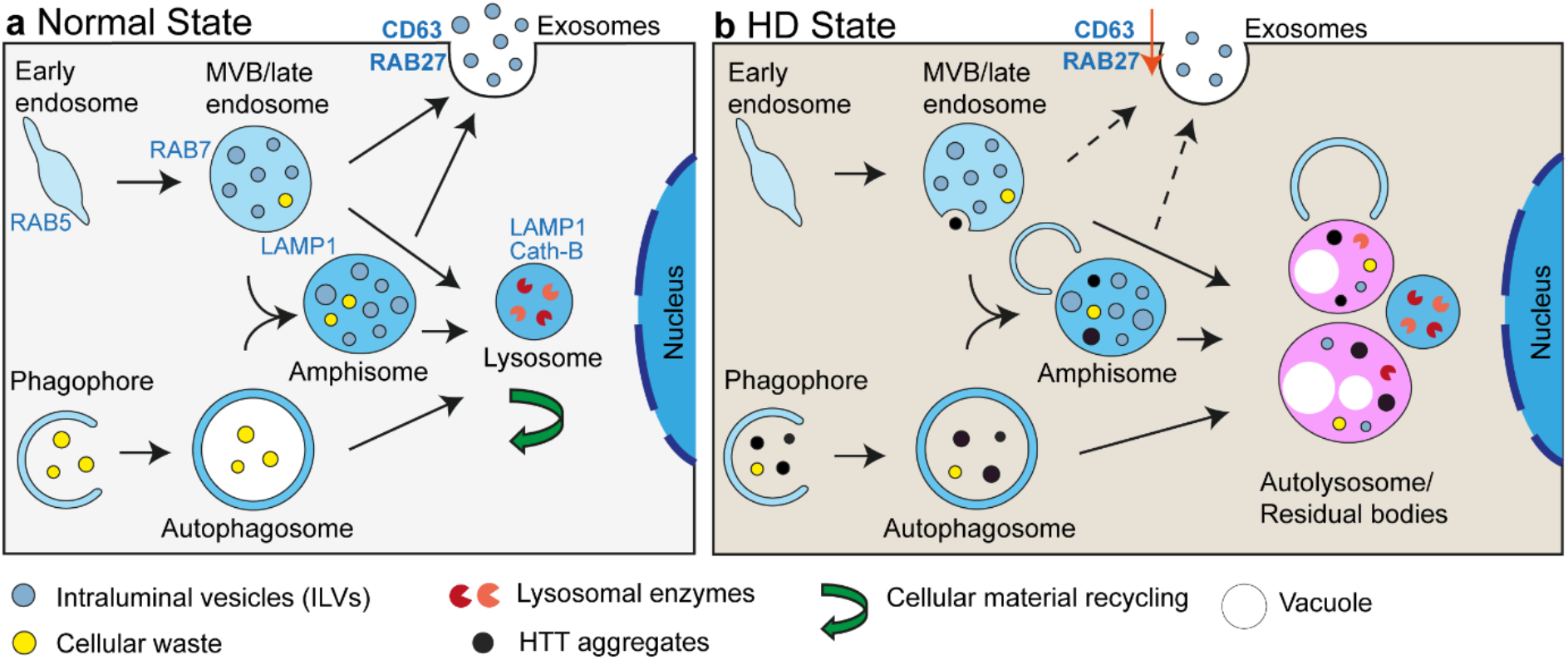
Schematic depiction of endolysosomal pathway dysregulation in Huntington’s disease. **a** The endolysosomal pathway under normal conditions. Early endosomes mature into multivesicular bodies (MVBs) containing many intraluminal vesicles (ILVs). MVBs can either merge with the plasma membrane to release ILVs as exosomes or undergo lysosomal degradation. Macroautophagy (autophagy) starts with the formation of a phagophore to engulf cellular contents. Expansion and maturation of the phagophore gives rise to the double-membraned autophagosome. Autophagosomes can either fuse with lysosomes to form an autolysosome or can merge with MVBs to form an amphisome. Amphisomes are believed to either undergo exocytosis or fuse with lysosomes. Eventually, MVBs, autophagosomes and amphisomes fuse with lysosomes and their contents are degraded and recycled. **b** In HD, it is likely that HTT aggregates are engulfed by a phagophore into an autophagosome and then enter amphisomes through the fusion of MVBs and autophagosomes to form this hybrid organelle. Alternatively, aggregates could be taken up by MVBs. Organelles over-loaded with HTT aggregates may become dysfunctional and/or be targeted for further autophagy and degradation. In this process, exocytosis through MVBs and amphisomes could be reduced and lysosomal or autophagic dysfunction may occur. This would result in the accumulation of MVBs/amphisomes and autolysosome/residual bodies containing HTT aggregates that were observed in HD brain sections.

We demonstrated by immunogold antibody labelling that mutant HTT is localized within the recruitment organelles, in electron lucent areas. Whilst no discernible fibrils or other structures were detected, the presence of small aggregates, well under 10 nm in diameter, cannot be excluded, and such nanostructures could act as polyQ recruitment centres. Why would aggregated HTT within the organelles recruit the exon 1 proteins, when fibrillar aggregates present at other locations in the cell, e.g. the nucleus, do not? It is possible that the acidic environment of these organelles has rendered aggregated HTT competent for HTT-exon1-43Q recruitment, possibly by stripping extraneous, non-specifically bound components from the growing end, or by inducing a more disordered state in which the polyQ region is more accessible to the label. The aggregated HTT present in the organelles may have entered the endolysosomal pathway via autophagy, through the fusion of autophagosomes with MVBs to form amphisomes. Alternatively, aggregated HTT may have been incorporated into the lumen of MVBs/endosomes, as occasionally, we observed immunogold HTT associated with small vesicular structures next to the recruitment organelles, an observation that requires further investigation (Fig. 9b).

MVBs and the endosome-lysosome system have been previously implicated in HD pathogenesis. HTT antibodies have been shown to bind to MVBs in HD patient brains [64] and functional MVBs were required for the efficient clearance of HTT aggregates via autophagic degradation [14]. Transfection of clonal striatal rat cells with HTT constructs stimulated the endosomal–lysosomal pathway, endosomal tubulation and the number, size and morphological variability of autophagic vacuoles and lysosome-like bodies [28]. RAB5 has been reported to recruit HTT to early endosomes via huntingtin associated protein 40 (HAP40) and mutant HTT results in altered endosome motility and endocytic activity [54]. In addition, inhibition of RAB5 enhanced polyQ toxicity in cell and fly models, potentially identifying a novel role for RAB5 in the early stages of autophagosome formation [60]. Mutant HTT impairs vesicle formation from recycling endosome by interfering with RAB11 activity [41], which is disrupted in a knockin mouse model of HD [40] and slows the recycling of the EAAC1 glutamate/cysteine transporter to the plasma membrane, leading to deficient glutathione synthesis and oxidative stress [42]. Our data are consistent with these reports in which total levels of RAB11 were not changed in HD [40].

We have demonstrated that mutant HTT dysregulates exocytosis *in vivo,* a novel mechanism through which mutant HTT disrupts the endolysosomal pathway (Fig. 9b). We found that the exosome proteins RAB27a and CD63 were both decreased by up to 30% in hippocampal lysates from zQ175 mice aged 6 months and R6/2 mice aged 8 and 12 weeks (Fig 8), indicating that the depletion of these proteins occurs throughout the entire disease course in these mouse models. CD63 is a tetraspanin that is enriched on the ILVs that are secreted as exosomes [58], and RAB27 functions in MVB docking at the plasma membrane [53]. These changes could reflect a shift in endolysosomal homeostasis away from exocytosis toward lysosome fusion and degradation, in response to the need to clear the chronically aggregating mutant HTT (Fig. 9b). In support of this, we detected an increase in cathepsin B in hippocampal lysates form 8- and 12-week-old R6/2 mice (Fig. 8c-f). Mutant HTT might induce a block in the pathway at the MVB / amphisome / autolysosome stages, a hypothesis supported by the frequently observed phagophore-like structures that appear poised to engulf the recruitment organelles (Fig. 9b).

We have established a method to correlate fluorescence microscopy images with their underlying EM-resolved structures on the same brain sections from mouse models of HD. In the work presented here, the fluorescent signal resulted from the recruitment of labelled exon 1 HTT proteins. This allowed us to determine the ultrastructure of a subset of the puncta seen by antibody staining and confocal microscopy. We have now adapted CLEM for use with fluorescently labelled antibodies on tissue sections to determine whether other aggregated HTT-rich puncta might contain other HTT aggregated states, or more complex structures as have been discovered for Lewy bodies in Parkinson’s disease [69,37,45]. Causative and risk-associated mutations, for a broad spectrum of neurodegenerative diseases, have been identified in many genes involved in the endolysosomal and autophagic pathways [46]. Our demonstration that proteins important for MVB-related exocytosis are considerably depleted throughout the disease course *in vivo* provides further evidence demonstrating that the endolysosomal pathway is dysregulated in HD and is likely to contribute to the pathogenic process.

## Author contribution

Conceptualization: Y.Z., G.P.B. and H.R.S.; Development of methodology: Y.Z. and T.R.P.; Sample preparation: Y.Z., T.R.P., C.L., J.B.W., K.S., E.J.S. and R.W.; Data Acquisition, Y.Z., C.L., K.S. E.J.S. and S.C.; Analysis: Y.Z. and T.R.P.; Writing original draft: Y.Z., G.P.B. and H.R.S.; Writing – review and editing: T.R.P, C. L., J.B.W., K.S., E.J.S., S.C., R.W., H.A.L., H.R.S.; Visualization: Y. Z.; Supervision: G.P.B. and H.R.S.; Funding Acquisition G.P.B. and H.R.S.

## Acknowledgements

We thank Natalya Lukoyanova for help with the EM data collection and comment on the manuscript, Alex Osmond, Lucy Collinson and Christopher Peddie for helpful discussions, David Houldershaw for computing support and Mark Turmaine for preparing osmium treated zQ175 and R6/2 EM samples as a comparison for CLEM data for this study. This work was supported by grants from the CHDI Foundation, Wellcome Trust awards 106249/Z/14/Z and 079605/2/06/2 and the UK Dementia Research Institute, which receives its funding from DRI Ltd, funded by the UK Medical Research Council, Alzheimer’s Society and Alzheimer’s Research UK.

