## Supplementary figures and video legends for "Correlative light and electron microscopy reveals that mutant huntingtin dysregulates the endolysosomal pathway in presymptomatic Huntington’s disease"

**Supplementary Table 1 Overview of the antibodies used in the present study**

| Antibody | Species | Dilution | | | Source | Product number |
|---|---|---|---|---|---|---|
| | | IF/IHC | WB | TEM | | |
| S830 | Sheep polyclonal | 1:1000 | 1:2000 | 1:20 | [1] | N/A |
| mEM48 | Mouse monoclonal | 1:100 | - | 1:20 | EMD Millipore | Cat# MAB5374 RRID: AB_10055116 |
| RAB5 | Rabbit polyclonal | 5 µg/ml | 1:1000 | - | Abcam | Cat# ab18211 RRID: AB_470264 |
| RAB7 | Rabbit monoclonal | 1:100 | 1:1000 | - | Abcam | Cat# ab137029 RRID: AB_2629474 |
| RAB11 | Mouse | - | 1:1000 | - | BD Biosciences | Cat# BD 610656 RRID: AB_397983 |
| RAB27A | Mouse monoclonal | 1:100 | 1:500 | - | Abcam | Cat# ab55667 RRID: AB_945112 |
| CD63 | Rabbit monoclonal | 1:100 | 1:1000 | - | Abcam | Cat# ab217345 RRID: AB_2754982 |
| LAMP1 | Rat monoclonal | 1:100 | 1:500 | - | Abcam | Cat# ab25245 RRID: AB_449893 |
| Cathepsin B | Rabbit monoclonal | 1:200 | 1:1000 | - | Abcam | Cat# ab214428 RRID: AB_2848144 |
| Cathepsin D | Rabbit monoclonal | 1:200 | 1:1000 | - | Abcam | Cat# ab75852 RRID: AB_1523267 |
| M6PR | Rabbit monoclonal | 1 µg/ml | 1:1000 | - | Abcam | Cat# ab124767 RRID: AB_10974087 |
| ATP5B | Mouse monoclonal | - | 1:25000 | - | Abcam | Cat# ab14730 RRID: AB_301438 |
| Rat Alexa Fluor 594 | Donkey polyclonal | 1:1000 | | | Thermo Fisher Scientific | Cat# A-21209 RRID: AB_2535795 |
| Sheep Alexa Fluor 647 | Donkey polyclonal | 1:1000 | | | Thermo Fisher Scientific | Cat# A-21448 RRID: AB_2535865 |
| Mouse Alexa Fluor Plus 555 | Donkey polyclonal | 1:1000 | | | Thermo Fisher Scientific | Cat: A32773 RRID: AB_2762848 |
| Goat Alexa Fluor 594 | Donkey polyclonal | 1:1000 | | | Thermo Fisher Scientific | Cat# A-11058 RRID: AB_2534105 |
| Rabbit Alexa Fluor Plus 555 | Donkey polyclonal | 1:1000 | | | Thermo Fisher Scientific | Cat# A32794 RRID: AB_2762834 |
| Mouse 20nm gold conjugated | Goat polyclonal | | | 1:50 | Abcam | Cat# ab27242 RRID: AB_954469 |
| Sheep 10nm gold conjugated | Rabbit polyclonal | | | 1:50 | Abcam | Cat# ab39609 RRID: AB_954445 |

IF = immunofluorescence; IHC = immunohistochemistry; WB = western blot; TEM = transmission electron microscopy.

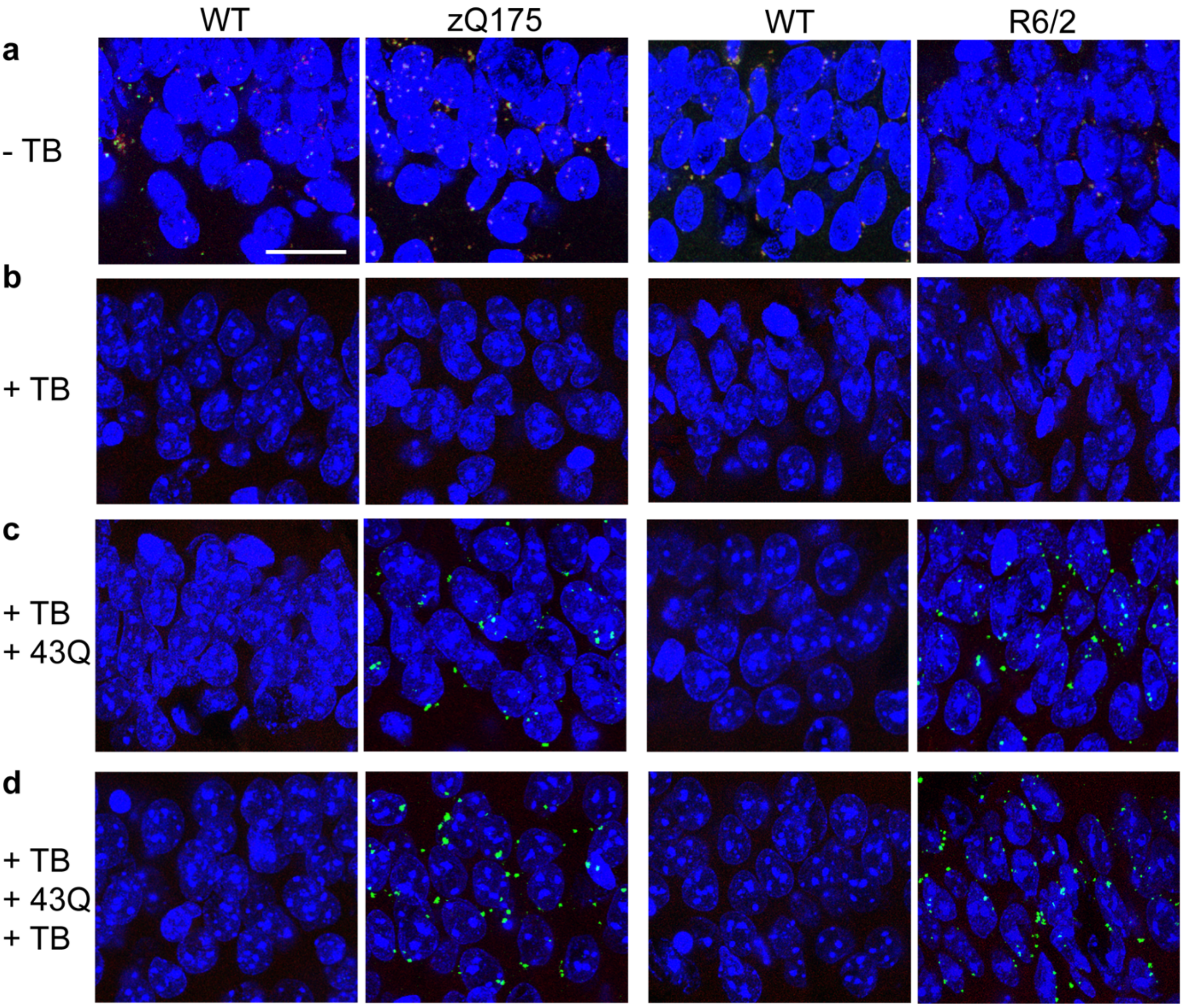


**Supplementary Fig. 1** Trueblack quenches brain tissue autofluorescence, but not the HTT-exon1-43Q fluorescence signal. **a** Without Trueblack treatment, autofluorescent puncta could be observed in brain sections from 6-month-old zQ175 and 12-week-old R6/2 mice and their wild-type littermates under both 488 nm and 561 nm lasers, using the filter settings for Alexa Fluor 488 and Alexa Fluor 568. The same imaging settings were used for all images in this figure, showing signals from both channels plus Hoechst for nuclear counterstaining. **b** After TrueBlack treatment, autofluorescent puncta were quenched in zQ175, R6/2 and wild-type brain sections. **c** When TrueBlack-treated zQ175, R6/2 and wild-type brains sections were incubated with the HTT-exon1-43Q-AF488 peptide, the recruitment signal was only observed on the zQ175 and R6/2 sections. **d** When Trueblack treated sections that had been incubated with the HTT-exon1-43Q peptide were again treated with TrueBlack, no quenching of the recruitment signal was observed. Scale bar: 20 µm. WT = wild-type, TB = Trueblack, 43Q = HTT-exon1-43Q-AF488.

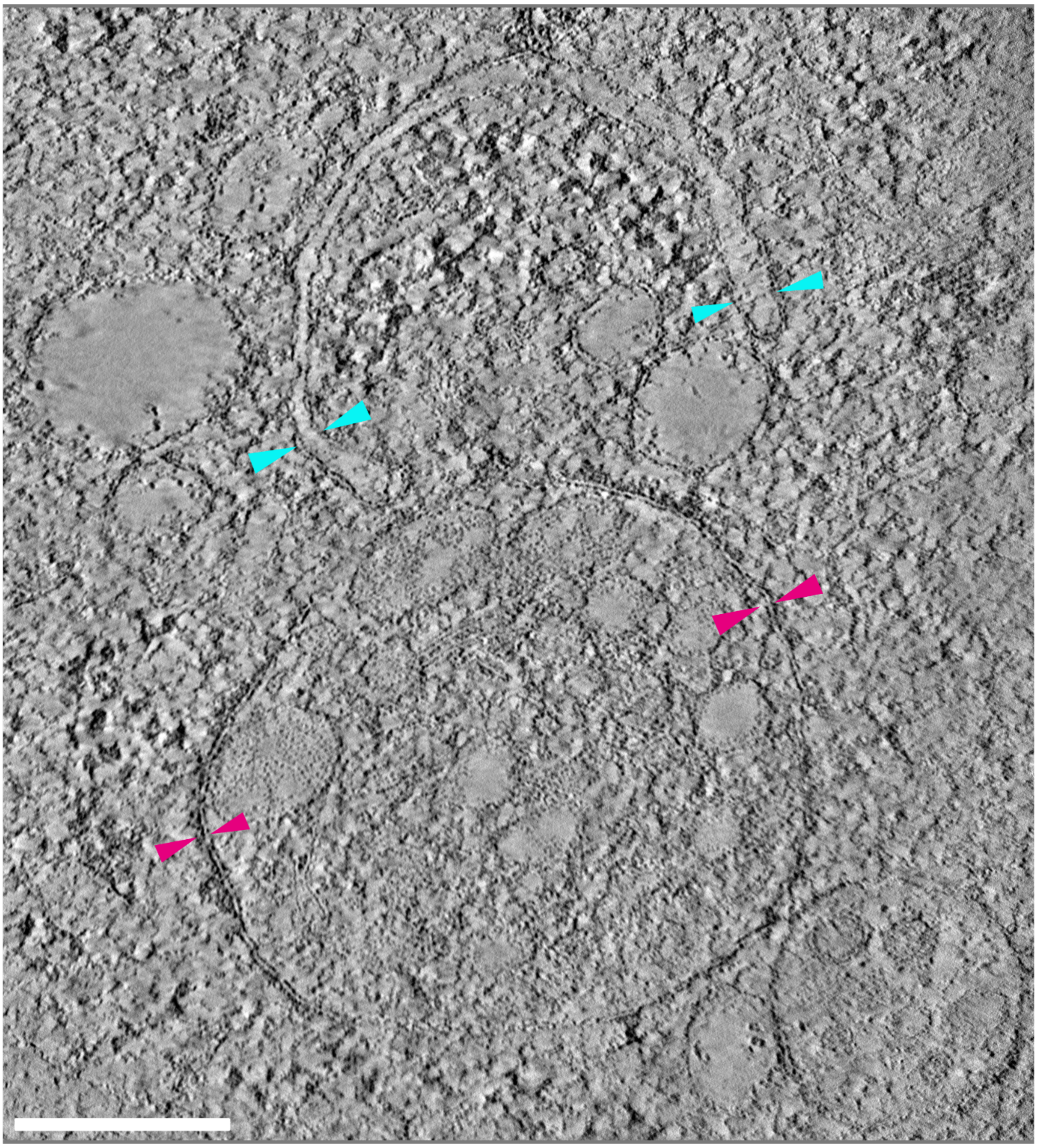

**Supplementary Fig. 2** Full-sized image of the organelle shown in Fig. 4g. Average of 10 central slices from the tomogram acquired for the organelle shown in Fig 4g. The single membrane of the recruitment organelle and double membrane of the nearby phagophore-like structure are indicated by magenta and cyan arrowheads, respectively. The lipid bilayers of the organelle and phagophore-like membranes are clearly visualized. Scale bar: 200 nm.

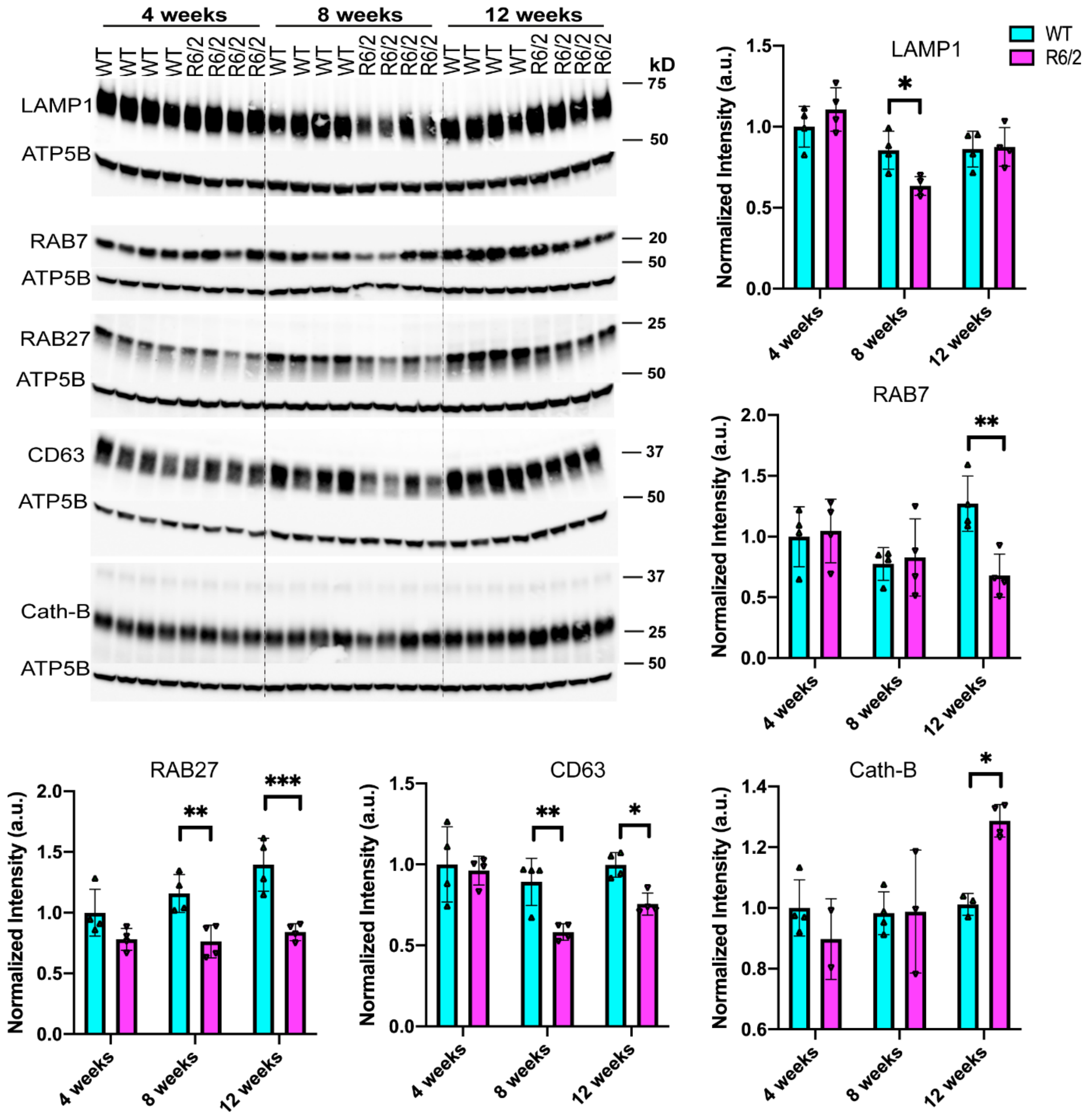


**Supplementary Fig. 3** Western blot of endolysosomal proteins in the R6/2 hippocampus. Western blotting of total protein hippocampal lysates from 4-, 8- and 12-week-old R6/2 mice and their wild-type littermates immunoprobed for either: LAMP1, RAB7, RAB27, CD63 or Cathepsin B (Cath-B) (n = 4 / genotype / age). ATP5B served as loading control. Quantification of the blot is shown on the right and below. The uncropped blots are shown in Supplementary Fig. 4, online resource 1. Statistical analysis was by two-way ANOVA. Data represented by mean ± S.D. *$p<0.05$; **$p < 0.01$; ***$p < 0.001$. Cath-B = Cathepsin B.

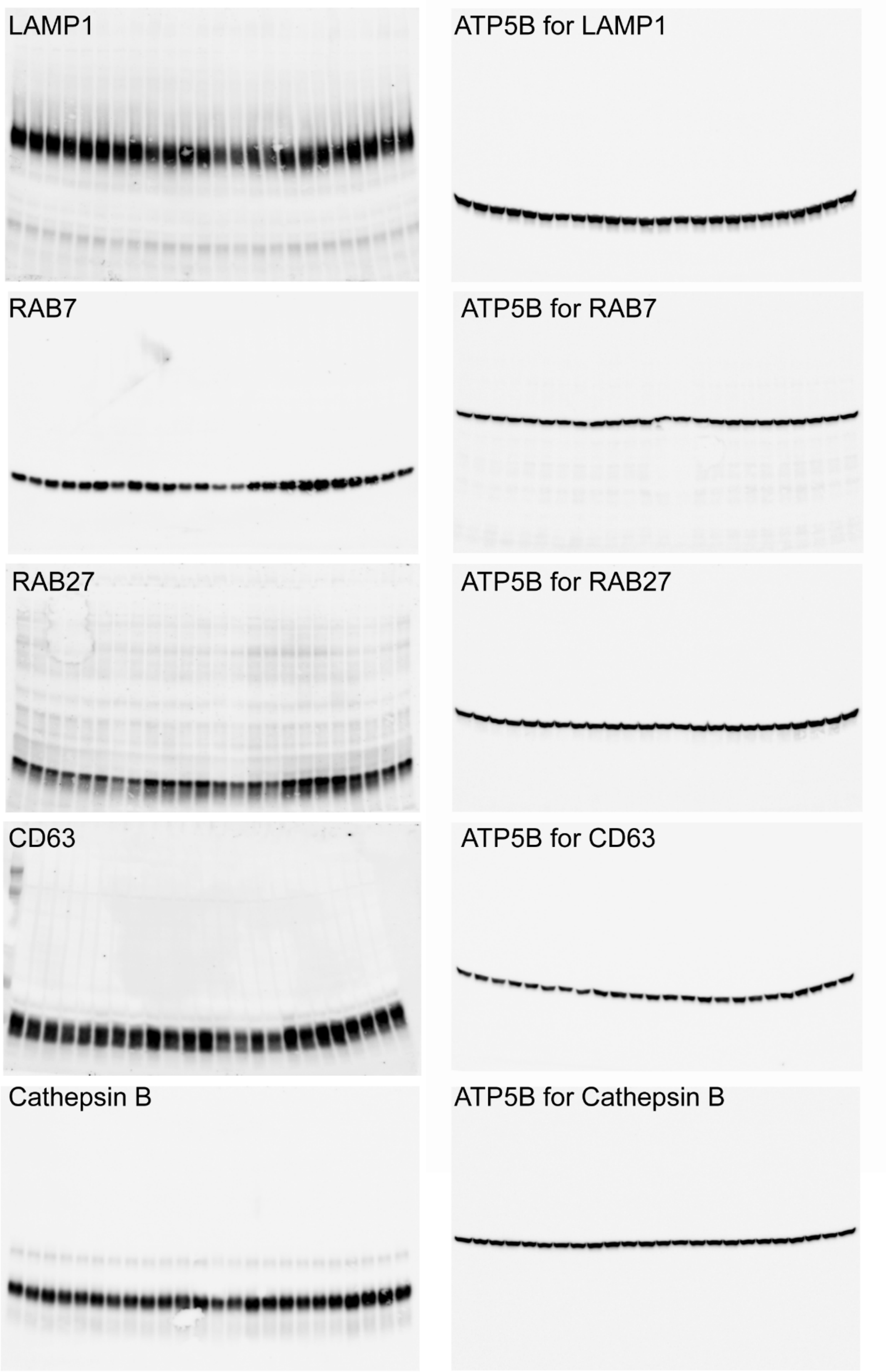


**Supplementary Fig. 4** Full-sized blots for the western blots presented in Supplementary Fig. 3.

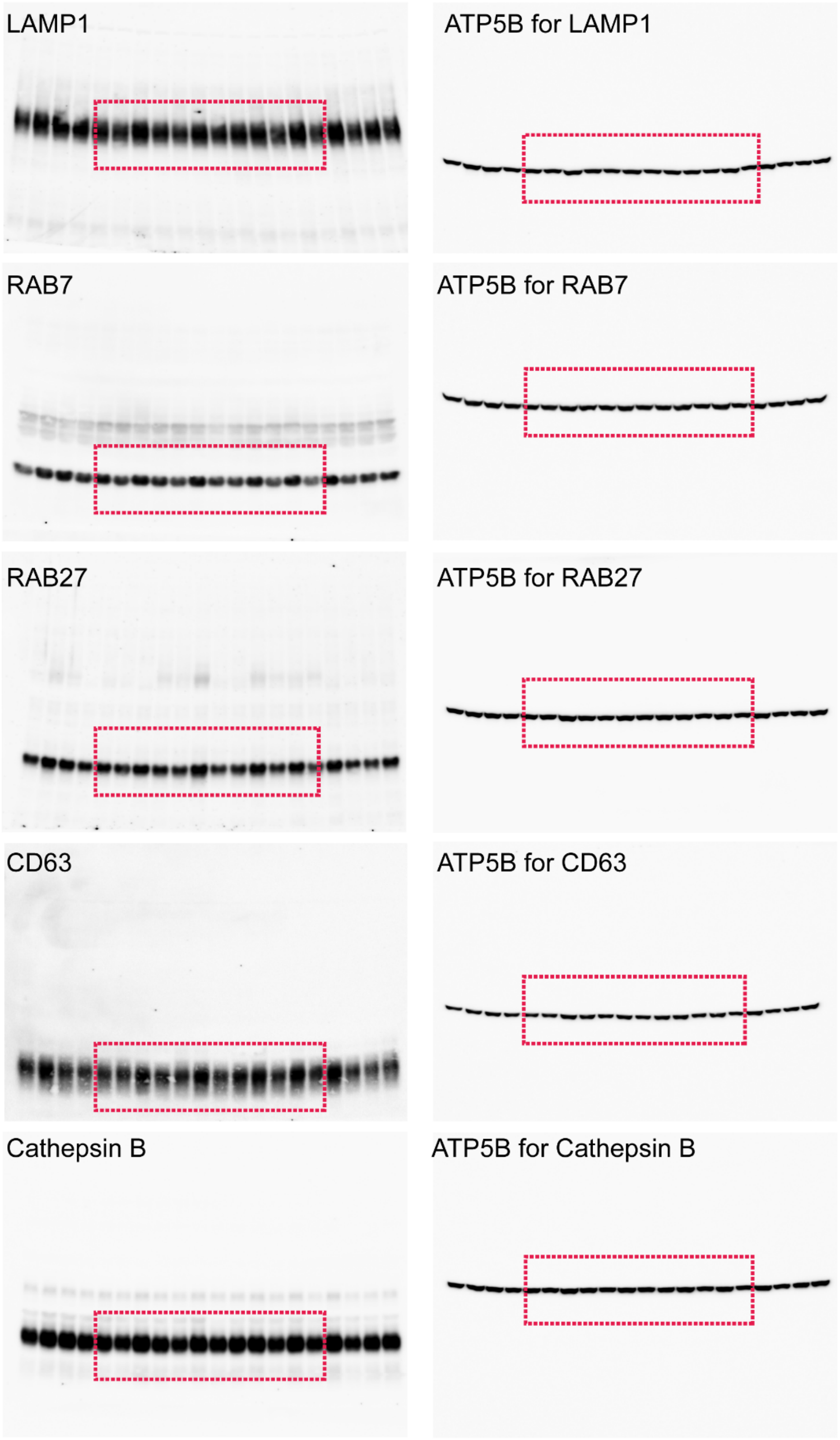


**Supplementary Fig. 5** Full-sized blots for the western blots presented in Fig. 8a. Red rectangular boxes indicate the areas shown in Fig. 8a.

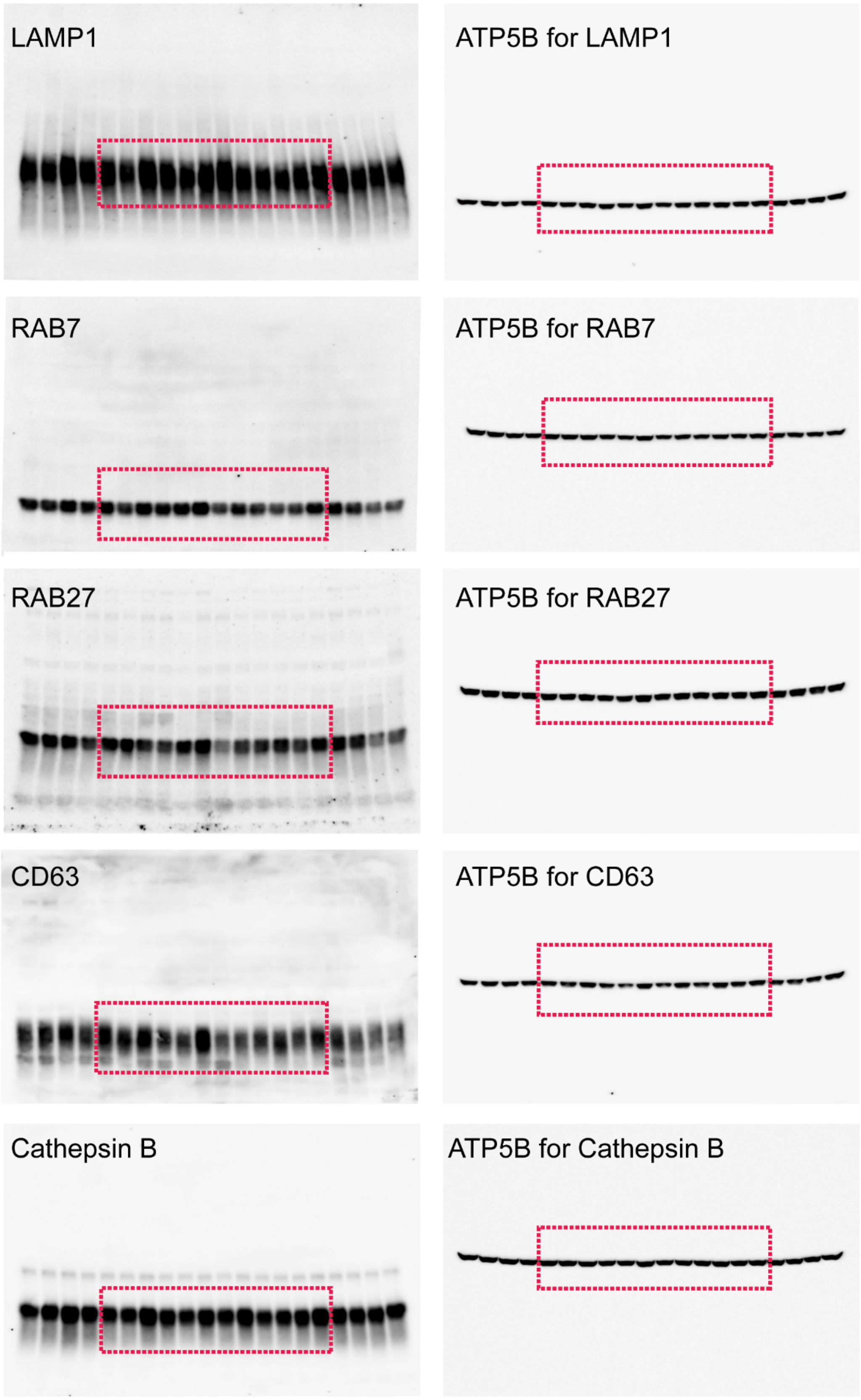


**Supplementary Fig. 6** Full-sized blots for the western blots presented in Fig. 8c. Red rectangular boxes indicate the areas shown in Fig. 8c.

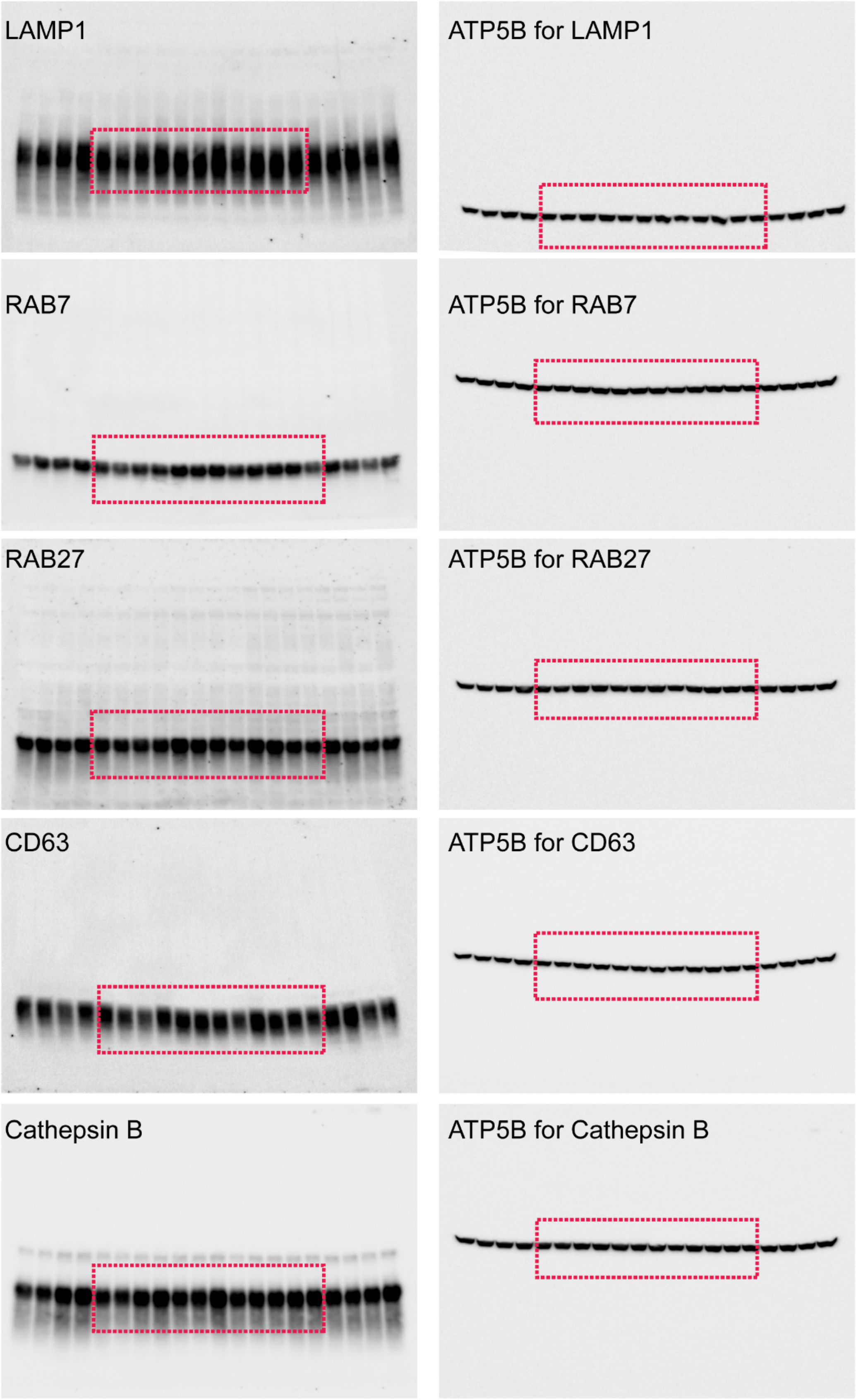


**Supplementary Fig. 7** Full-sized blots for the western blots presented in Fig. 8e. Red rectangular boxes indicate the areas shown in Fig. 8e.

### Supplementary Videos

**Supplementary Video 1, online resource 2** Tomogram related to Fig 4g, a recruitment site (shown in magenta in segmentation) identified by CLEM in 6-month-old zQ175 mice.

**Supplementary Video 2, online resource 3** Tomogram related to Fig. 6b, mutant HTT containing organelle resembles MVB/amphisome in 6-month-old zQ175 mice.

**Supplementary Video 3, online resource 4** Tomogram related to Fig. 6c, mutant HTT containing organelle resembles autolysosome in 6-month-old zQ175 mice. Intraluminal lamellar structures are shown in cyan in segmentation.

**Supplementary Video 4, online resource 5** Tomogram related to Fig. 6d, mutant HTT containing organelle resembles autolysosome/residual body in 6-month-old zQ175 mice.
